# Ubiquitous trophic niche partitioning drives species coexistence and biomass on coral reefs

**DOI:** 10.64898/2026.09.22.753627

**Authors:** Jordan M. Casey, Simon J. Brandl, Nina M. D. Schiettekatte, Samuel Degregori, Giovanni Strona, Alexandre Mercière, Fabien Morat, Mayalen Zubia, Christopher P. Meyer, Valeriano Parravicini

## Abstract

Coral reefs support an extraordinary diversity and biomass of fishes. However, whether species coexist through fine-scale niche partitioning, and whether this impacts ecosystem functioning, remain unresolved. We combined DNA metabarcoding and stable isotope analysis to reconstruct a high-resolution food web and quantify the trophic niches of 2,060 reef fishes across 261 species. We observed extreme trophic partitioning: species pairs diverged by a median of 93% (DNA metabarcoding) and 86% (stable isotopes), consistently overlapping far less than expected by chance. Further, communities with greater trophic complementarity supported higher standing-stock biomass, particularly at high species richness. Our results confirm limiting similarity and trophic complementarity as the driving forces of coexistence and ecosystem functioning in the ocean’s most diverse ecological community.

## Main Text

How so many species can coexist in diverse ecosystems is a central question in ecology, fueling decades of debate over the mechanisms of community assembly (*1–4*). These debates center around the relative importance of stochastic neutral *versus* deterministic niche-based processes. Neutral theory predicts that biodiversity patterns emerge solely based on demographic stochasticity and dispersal limitations (*5–7*), but this model may fail to explain patterns in complex ecosystems such as rainforests and coral reefs (*8*). In contrast, niche-based theory emphasizes resource use and niche partitioning as the core drivers of biodiversity patterns (*9*). At the core of niche-based theory lies the principle of limiting similarity, which hypothesizes that two species cannot coexist if they compete for the same resources (*4*). While this has been empirically demonstrated at small scales, especially among species pairs, many researchers have proposed that trophic redundancy may be inevitable in diverse, tropical animal communities, with species overlapping more in resource use as richness increases (*10*). However, these claims are often supported by coarse dietary datasets that reduce thousands of distinct prey items into a few simplified categories, which are then used to assign consumers to trophic groups (*11, 12*).

Tropical coral reefs are the ocean’s most diverse and productive ecosystem (*13*), making them an ideal system to test whether niche partitioning promotes species coexistence and biomass production in complex communities. Yet, their tremendous diversity has also made them fairly intractable, with multitudes of cryptic organisms and obscure trophic pathways (*14, 15*). Fishes represent the dominant consumer pathway on coral reefs, with diverse species assimilating energy and nutrients from countless primary and intermediary sources to produce a rich pool of standing stock biomass and oases of diversity (*16*). Oftentimes, hundreds of fish species co-occur on a single reef system, but our current understanding of coral reef food webs has remained surprisingly simplistic. To date, existing high-resolution dietary datasets that examine the trophic niches of consumers and use multiple, complementary techniques only include a few species (*17, 18*), or a single trophic guild (*19*). Consequently, no current empirical dataset allows us to test the drivers of coexistence in fishes at the community level. This also hampers our ability to predict and manage one of the primary services of coral reefs: the provision of harvestable reef fish biomass, which is a direct product of resource use and coexistence.

Behavioral observations and the visual examination of gut contents are the dominant tools used to map trophic interactions in reef fishes. However, especially in high-diversity systems, these approaches often overlook cryptic organisms and provide poor taxonomic resolution of partially digested prey items (*20*). Molecular approaches and biochemical techniques have revolutionized our discernment of trophic interactions (*19, 21*). DNA metabarcoding of gut contents or fecal samples provides exceptional resolution of prey items, while stable isotope ratios (δ^13^C and δ^15^N) reveal the long-term trophic preferences of an organism. Together, these approaches are highly complementary: metabarcoding captures a snapshot of unparalleled dietary resolution while the broad time window of stable isotope analysis guards against spatial and temporal biases associated with stochasticity in prey consumption (*22*).

To investigate whether niche-based processes govern species coexistence in the most diverse marine ecosystem, we quantified trophic partitioning across an entire coral reef fish community and tested its potential impacts on community assembly and biomass. To infer the trophic niches of coral reef fishes, we built fish gut content metabarcoding and stable isotope datasets of unprecedented scope and resolution, including 2,060 individuals across 261 species and 48 families of reef fishes from Mo’orea, French Polynesia, an oceanic island in the South Pacific (Fig. 1; Fig. S1). The fish consumers in our dataset span nearly two orders of magnitude (1.3 cm to 120 cm) in body size, capturing cryptobenthic (< 5 cm) to large, mobile fishes, and comprising approximately 86% of the total fish community biomass around Mo’orea (Fig. S2). These data uncover an unparalleled, high-resolution coral reef food web characterized by ubiquitous trophic niche partitioning. We pair this with a 19-year time series of fish communities around Mo’orea to reveal trophic complementarity as a key mechanism underpinning reef fish species richness and standing stock biomass, providing empirical evidence for a complementarity-driven link between biodiversity and ecosystem functioning on coral reefs. Our results highlight the overarching importance of niche-based processes in driving species coexistence and functioning of highly diverse, and often threatened, tropical ecosystems.

**Fig 1.**
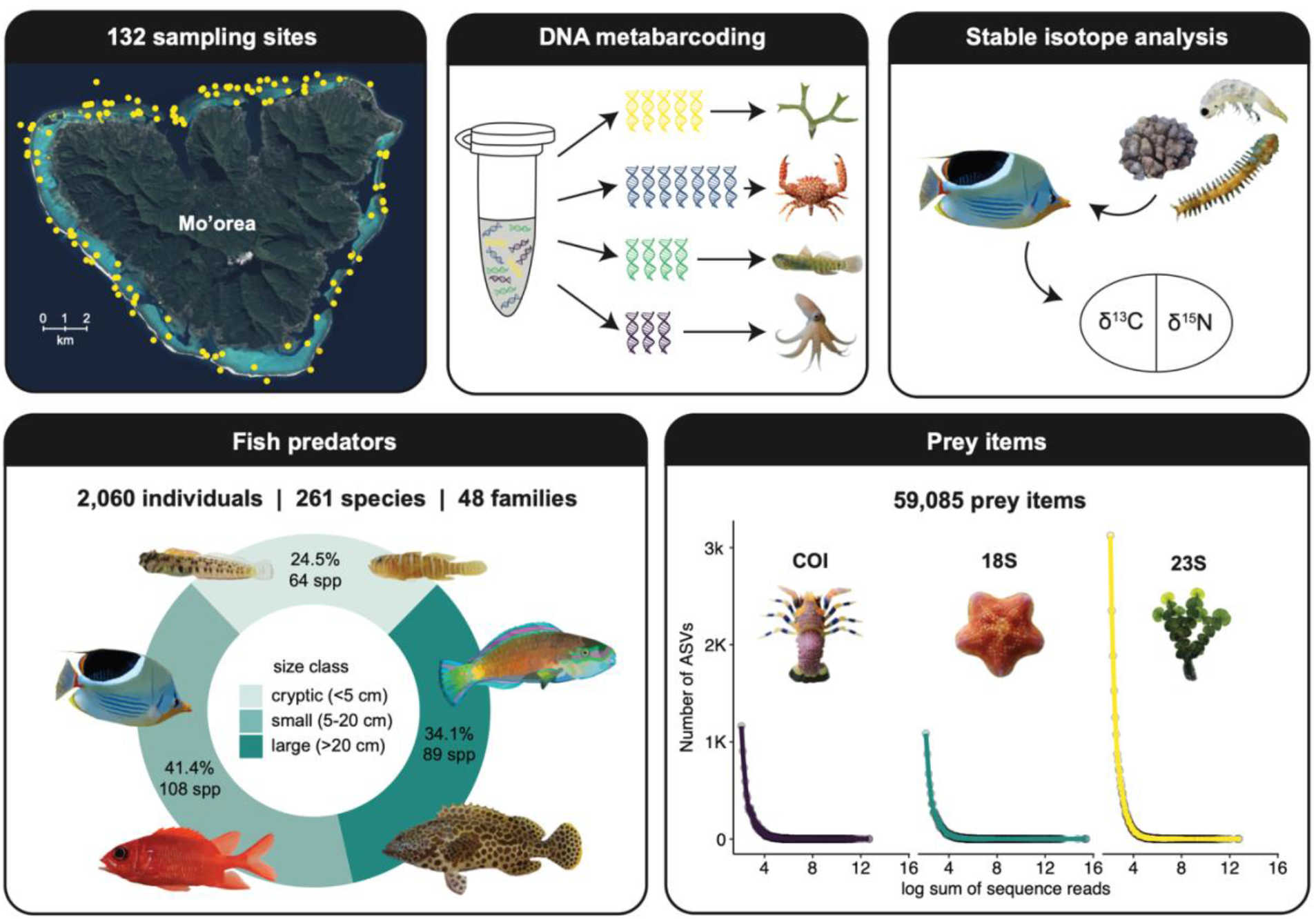
Sampling sites, analytical approaches, fish collections, and summary of prey items based on molecular data. Fish collections were conducted at 132 sites around Mo’orea, French Polynesia, with sites ranging from the lagoon to the reef crest. Fish diet and trophic niches were analyzed with molecular (DNA metabarcoding of fish gut contents) and biochemical (stable isotope analysis of fish muscle tissue) approaches. Fish collections included 2,060 individuals across 261 species and 48 families, covering a large range of fish size classes (1.7 cm to 1,120 cm). With three molecular markers (COI, 18S, and 23S), DNA metabarcoding of fish gut contents identified 59,085 prey items based on amplicon sequence variants (ASVs).

### Extreme trophic niche partitioning across a coral reef food web

In our DNA metabarcoding dataset of gut contents, after strict data filtering, we generated 52,215,064 sequence reads across three markers: 9,073,386 for COI, 35,090,517 for 18S, and 8,051,161 for 23S (Fig. 1). Sequence reads were grouped by amplicon sequence variants (ASVs), which are distinct genetic signatures and an approximation for prey items. We identified 59,085 ASVs: 15,062 for COI, 13,776 for 18S, and 30,247 for 23S. We created a single binary dataset of presence-absence across the three markers, merged by the lowest robust taxonomic identity, which yielded 2,258 taxonomically unique prey items. This approach merges prey items at their lowest taxonomic level and does not consider relative abundances of prey items, thus providing a highly conservative framework to estimate niche partitioning among coral reef fishes.

Nonetheless, this simplified, merged data frame reveals a remarkably complex network of fish consumers and the wide range of prey items that they consume, including many unanticipated trophic pathways and representing a nearly complete fish-centric coral reef food web (Fig. 2A).

**Fig 2.**
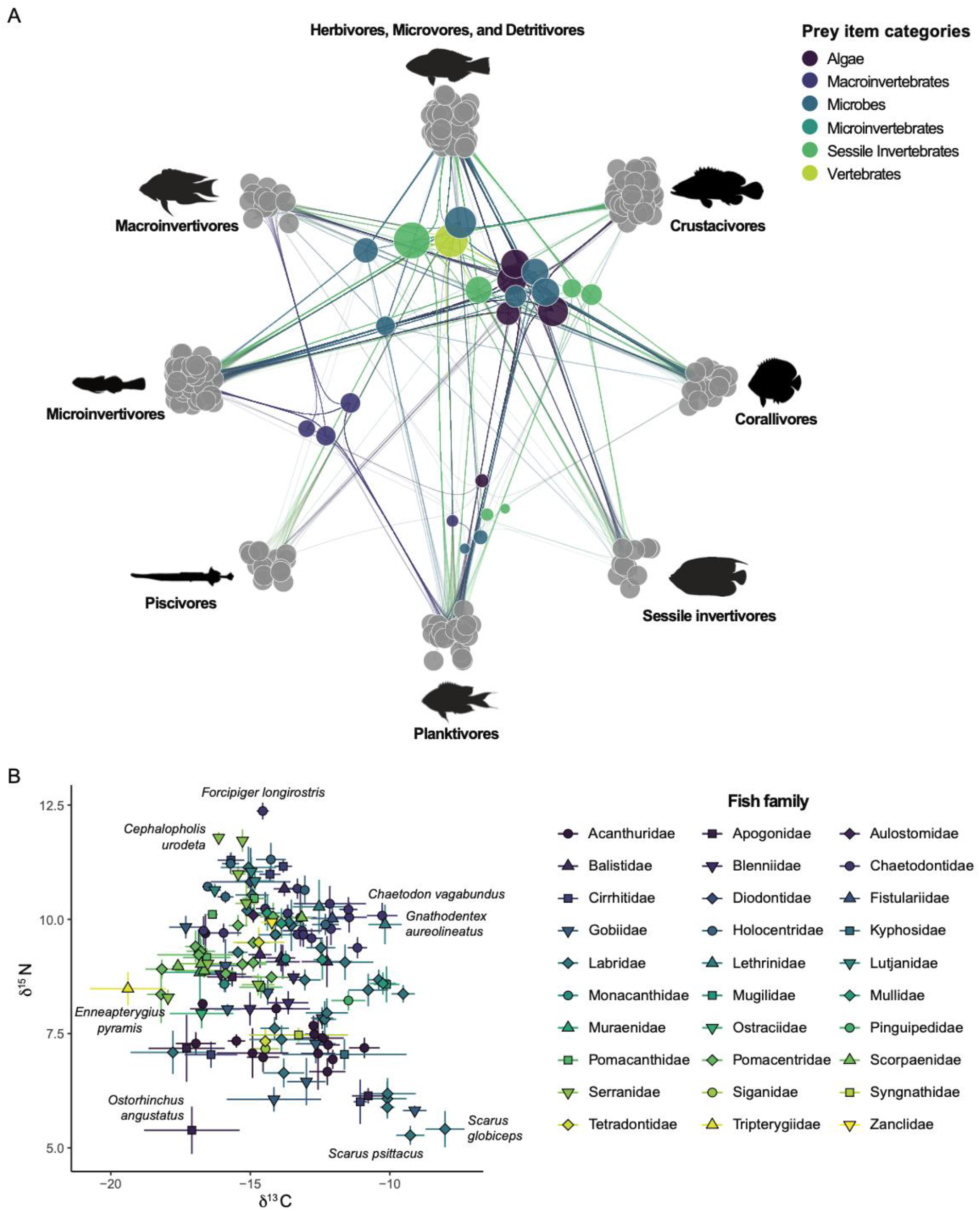
The complexity and diversity of trophic pathways across a reef fish food web. (**A**) Aggregated trophic network plot based on DNA metabarcoding data. (**B**) Mean δ^13^C and δ^15^N (± standard error) of reef fish species, grouped by fish family, based on bulk stable isotope data. Species are only included if more than three individuals were sampled. Species names are included for species that occupy the outer vertices of isotopic niche space.

We bolstered our metabarcoding dataset with stable isotope analysis on a subset of 216 fish species. Fishes were broadly distributed across δ^13^C-δ^15^N niche space, revealing substantial variation in resource use and trophic position (Fig 2B). Scarine labrids (parrotfishes), which primarily consume basal resources such as algae and detritus (*23*), showed low δ^15^N and enriched δ^13^C values, whereas *Forcipiger longirostris*, a caridean shrimp specialist (*24*), occupied the highest trophic position, highlighting unexpected pathways supporting mid-tier consumers on coral reefs (*25*). Cryptobenthic fishes, which comprised 24.7% of sampled species but are often absent from food web studies, occupied distinct isotopic positions, including *Enneapterygius pyramis*, whose carbon signature was consistent with reliance on pelagic sources and thus a pathway to vector pelagic production into reef food webs (*26, 27*). Their inclusion is therefore critical for resolving food web structure given their disproportionate contribution to biomass production through rapid growth and turnover (*28, 29*).

To quantify trophic niche partitioning across fish species with dietary metabarcoding and stable isotope data, we measured niche segregation for each pair of species (see Materials and Methods). From the dietary metabarcoding data, we demonstrate a striking degree of trophic niche partitioning among fish species, with a median of 93% (average of 89.7%) niche segregation across all species. Within families, fish species had a median of 81.6% niche segregation, and all species had at least 31.6% unique niche space (lowest niche segregation: *Callogobius* sp. and *Paragobiodon lacunicolus*; Fig. 3A). Within genera, species had a median of 73.7% niche segregation, and all species had at least 33.2% niche segregation (lowest niche segregation: *Callogobius sclateri* and *Callogobius* sp.; Fig. S3A). Similarly for the isotopic data, we observed strong niche partitioning, with a median of 86.2% (average of 82.2%) niche segregation across all species. Within families, fish species had a median of 74.8% niche segregation, and within genera, species had a median of 66.7% niche segregation (Fig. 3B). Within genera and families, all species had at least 31.5% niche segregation (lowest niche segregation: *Melichthys niger* and *Melichthys vidua*; Fig. S3B).

**Fig 3.**
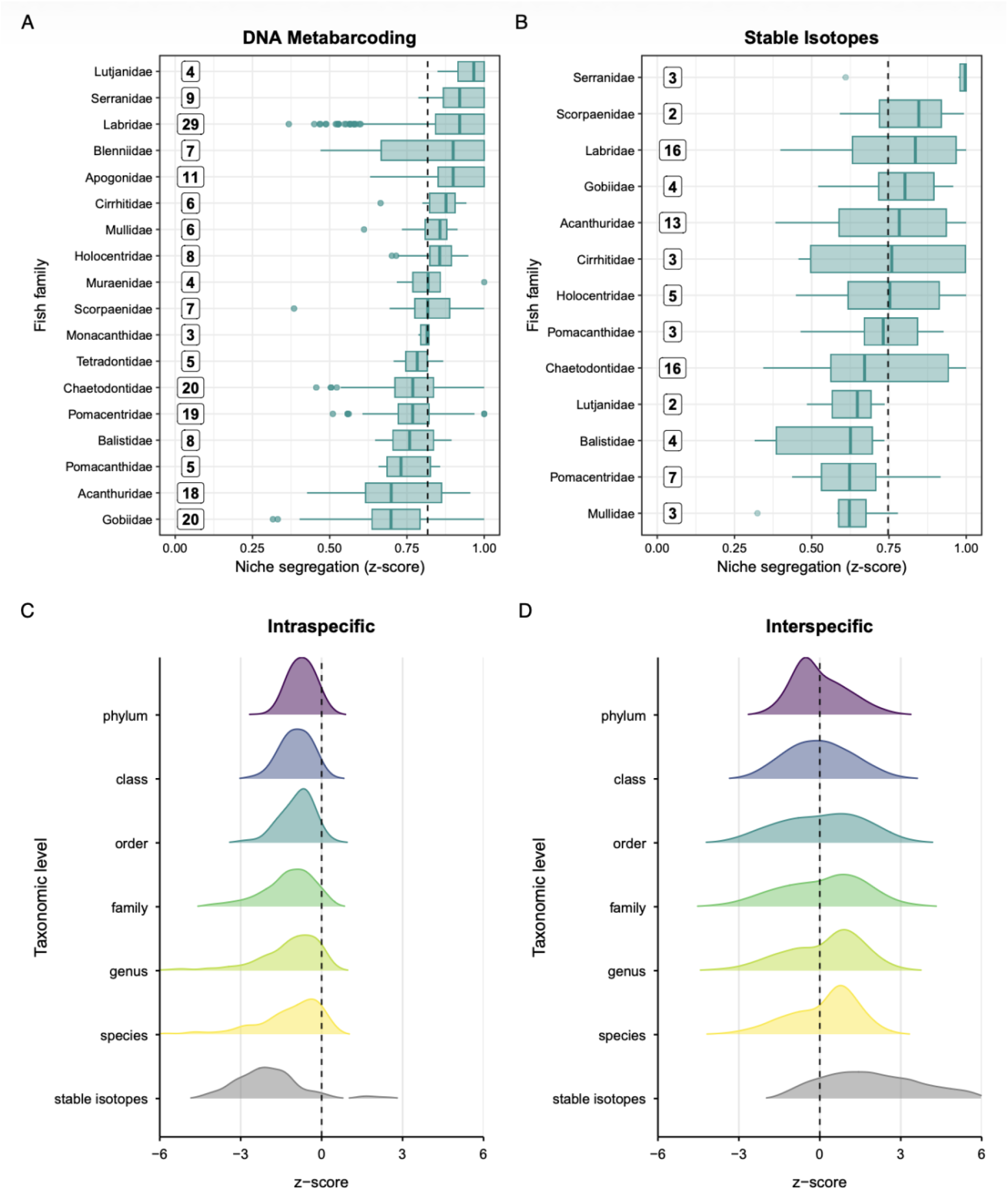
Trophic niche partitioning across fish species based on molecular and isotopic approaches. (**A** to **B**) Percent niche segregation (z-scores) between species pairs, within fish families, based on (**A**) Pianka overlap for molecular and (**B**) Euclidean distances for isotopic data. The numbers to the left of the boxplots indicate the total number of species sampled within that family. The boxplots represent the medians and interquartile ranges, and the whiskers represent 1.5 times the interquartile range. Points outside the boxplot whiskers represent outliers. Fish families are ranked from highest (top) to lowest (bottom) trophic niche segregation based on median values. The dashed vertical lines represent the average niche segregation of all species across the (**A**) 18 fish families in the molecular dataset and (**B**) 13 fish families in the isotopic dataset. (**C** to **D**) Likelihood of trophic niche overlap (z-scores) in observed molecular and isotopic data as compared to a null model between (**C**) intraspecific (individuals within species) and (**D**) interspecific (between species) species pairs. For the molecular data, trophic niche overlap is examined at multiple taxonomic levels, from phylum to species, for each prey item. When intraspecific or interspecific overlap is to the right of the dashed vertical line at zero, there is less overlap than expected by chance.

For the dietary metabarcoding data, across families, Lutjanidae (snappers, N=4 species) had the highest median niche segregation (96.6% ± 1.7 SE), while Gobiidae (gobies, N=20) had the lowest (69.8% ± 0.7 SE; Fig. 3A). For the isotopic data, across families, Serranidae (groupers, N=3 species), had the highest median niche segregation (99.6% ± 6.4 SE), while Mullidae (goatfishes, N=3), had the lowest median niche segregation (62.2% ± 6.3 SE; Fig. 3B).

Generally, mesopredators, such as wrasses, snappers, and groupers exhibited the highest unique niche space, while herbivorous and detritivorous fishes, such as parrotfishes and surgeonfishes, displayed the lowest unique niche space. These findings track patterns of consumption rates and resource availability across coral reef fishes: predatory fishes are more likely to eat periodic, large meals, resulting in the inflation of dietary uniqueness across individuals, while herbivorous fishes feed throughout the day on more homogenous, immobile resources to achieve sufficient nutrient intake, leading to a higher likelihood of dietary overlap in species pairs (*30*). Yet, remarkable dietary partitioning existed even among closely-related herbivorous and detritivorous fishes, which mirrors dietary differences revealed by DNA metabarcoding of fecal matter from large herbivorous mammals in African savannas (*31, 32*) and a subset coral reef herbivores in Hawaii (*33*). Overall, we demonstrate extremely limited trophic redundancy in coral reef fishes, upholding foundational ecological theory: even in one of the most diverse assemblages of vertebrates on our planet, every species has a largely distinct trophic niche, consistent with limiting similarity as a basis for stable coexistence. Notably, our findings are conservative because they do not account for fish foraging across space (e.g., microhabitat use) or time (e.g., noctural *versus* diurnal), two common niche axes that contribute to trophic partitioning, especially within herbivores (*34, 35*).

To examine the prevalence of niche partitioning, we evaluated niche segregation within and across species against a null model (Fig. 3C-D). Across all prey item taxonomic ranks (phylum to species), we detected notable intraspecific trophic niche overlap, indicating high dietary similarity among individuals within species, particularly when prey items are defined to the genus and species level. In stark contrast, overlap among interspecific pairs progressively declined as prey were assigned to the rank of order, family, genus and species. Indeed, at the genus and species levels, 60.1% and 62.3% of species pairs, respectively, showed less trophic niche overlap than expected by chance, demonstrating weak interspecific competition for prey. Given the stochasticity of prey ingestion and individual dietary heterogeneity, gut content metabarcoding might be expected to overestimate niche segregation because it only captures a snapshot of fish diet. Yet, our molecular results were corroborated by the isotopic data, which integrates dietary sources over space and time: 85.6% of species pairs exhibited less trophic niche overlap than expected by chance (Fig. 3D). The stronger niche segregation detected by time-integrated isotopic data suggests that short-term dietary stochasticity, driven by fluctuating prey availability, may inflate niche overlap captured by dietary metabarcoding at a single time point. Thus, rather than an overestimation, DNA metabarcoding may provide a more conservative estimate of niche segregation.

Overall, both methods demonstrate a clear pattern: coral reef fishes exhibit extreme trophic niche partitioning, and current estimates of trophic redundancy vastly underestimate dietary uniqueness of reef fishes. By reducing reef fish species to broad functional groups or trophic guilds, we are overlooking biological diversity and the irreplaceable contribution of each fish species to ecosystem functioning. By extension, coral reef food webs are far more complex than previously estimated, and the tremendous diversity of potential prey items on reefs gives rise to a similarly diverse range of specialized, trophic niches. The occurrence of 139,122 trophic interactions in our food web provides a holistic, yet conservative, glimpse into the multitude of interactions occurring on a tropical coral reef. In the grand scheme of biodiversity across coral reefs, the reefs around Mo’orea, French Polynesia are less diverse than those in the Western Pacific, so the complexity of coral reef food webs in the center of ocean diversity, such as the Coral Triangle (*36*), would likely be even more astounding.

Modern coexistence theory posits that species coexistence is stabilized when intraspecific competition for resources outweighs interspecific competition (*37*). Our results clearly reveal the stabilizing dynamics of intra- *versus* interspecific competition that underlie coexistence in reef fishes. Detecting this signal required moving beyond the broad trophic and taxonomic categories that consumers and their prey are often lumped into (e.g. herbivores or invertivores; crustaceans or worms), which are insufficient to capture the complexity of coral reef trophic interactions and their ecological consequences (*20*). Researchers have attempted to piece together holistic food webs (*38, 39*); however, the taxonomic resolution of prey items is often limited, which impairs our ability to evaluate the ecosystem level consequences of losing fish species in response to anthropogenic stressors (*40*). High-resolution dietary data could also reveal whether species compensate for competitor loss by broadening their trophic niche, evaluating complementarity as a buffer against biodiversity loss (*41*).

### The role of dietary complementarity for community assembly and biomass

Biodiversity is frequently invoked as a driver of ecosystem functioning through functional complementarity (*42*). Our data offer a unique opportunity to examine how extreme niche partitioning impacts community assembly and biomass in a highly diverse system. Using 19- years of fish community data from Mo’orea paired with our dietary metabarcoding data, we quantified community-level niche segregation across species pairs compared against a null model, using unweighted and biomass-weighted metrics (see Materials and Methods).

The two metrics revealed contrasting patterns. Unweighted segregation was centered near zero (Fig. 4A), indicating that local assemblages are neither more nor less segregated than expected from the regional species pool. Thus, based on presence-absence alone, extreme niche partitioning is simply mirrored at the local scale. In contrast, biomass-weighted segregation was consistently negative, indicating greater trophic overlap than expected by chance among the most dominant consumers (Fig. 4D). Crucially, the relationships with biomass also diverged. High unweighted segregation was associated with lower standing stock biomass (Fig. 4C), while biomass-weighted segregation showed the opposite pattern: standing stock biomass increased with greater niche segregation of the most dominant consumers (Fig. 4F). Thus, biomass peaked in species-rich assemblages where the dominant, high-biomass species show low trophic niche overlap (Fig. 4E).

**Fig 4.**
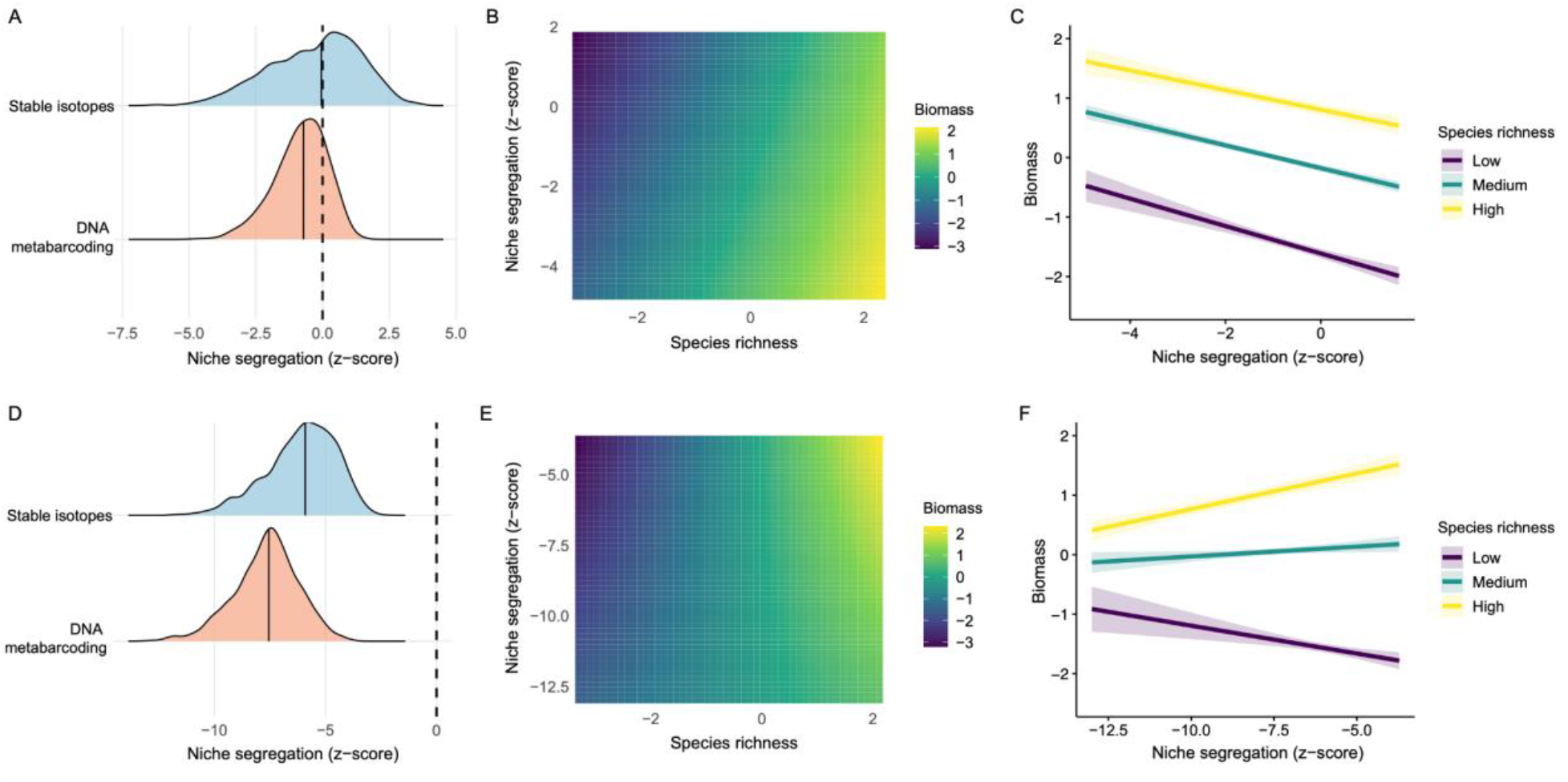
The effects of species richness and trophic niche partitioning on standing stock biomass across reef fish communities. Community-level trophic niche segregation was quantified across species pairs using (**A** to **C**) unweighted and (**D** to **F**) biomass-weighted Pianka-based metrics. (**A, D**) The distribution of community-level niche segregation (z-scores) across transects relative to null models that reshuffles species identities and biomasses across regional pools while retaining species-level trophic niches. Positive z-scores indicate greater niche segregation than expected by chance. Densities are shown separately for molecular and isotopic data. (**B, E**) The interactive effects of species richness and niche segregation on standing-stock biomass. Color indicates the posterior predicted biomass on the log scale. (**C, F**) The relationship between niche segregation and biomass at low, medium, and high species richness (minimum, median, and maximum observed). (**C**) Biomass declines with higher unweighted niche segregation but (**F**) increases with biomass-weighted segregation in high diversity communities.

The ecological consequences of trophic niche complementarity thus depends on which species contribute to community biomass. Rare specialists can generate substantial niche partitioning without significantly contributing to community standing stock biomass, while fine-scale niche partitioning among dominant consumers is linked to greater biomass. This likely reflects competition for locally abundant resources: dominant consumers overlap more than expected when a common resource supports much of the community, but communities with higher trophic niche partitioning can sustain greater biomass. Thus, biodiversity-biomass relationships may depend less on the total extent of niche partitioning than on how complementarity is distributed among dominant species that channel energy through the ecosystem. This finding extends previous work linking herbivore consumer diversity with functional complementarity and ecosystem functioning on coral reefs (*43*); previously demonstrated in a two-species cage experiment, we show this across a diverse, natural community of reef fishes.

The challenges of quantifying interactions in hyperdiverse food webs have led most experiments and simulations of biodiversity-ecosystem functioning relationships to rely on relatively simple, low diversity fractions of communities (*44, 45*), limiting our understanding of how processes scale to ecosystems with hundreds of species. In turn, large-scale, survey datasets can reveal correlative relationships between species richness and functioning, but they provide little insight into the underlying mechanisms (*15*). On coral reefs, large-scale surveys link species richness and functional diversity with biomass (*46, 47*), but our results suggest that coarse functional assignments in these studies can mask the effects of trophic complementarity on functioning. On the other hand, claims of functional redundancy and extreme trophic versatility among reef fish species have led to the assertion that species diversity has a limited impact on key ecosystem processes on coral reefs (*48*). Similarly, high diversity reef systems have been characterized as redundant, with ecologically equivalent species collapsing into a limited set of functional entities (*11*). Our results challenge these views, revealing that the unique trophic space occupied by hundreds of reef fish species can support stable coexistence and high standing biomass, a key ecosystem service that supports millions of people that depend on coral reefs worldwide.

Across other ecosystems, trophic complementarity has been linked to standing stock biomass and biomass production (*49, 50*). Yet, this has never been validated across a large, empirical dataset. Moreover, most empirical studies have focused on lower trophic levels, particularly simple invertebrate-plant interactions that are amenable to field-based experimental manipulations (*51*). Our high-resolution dataset provides an opportunity to extend a core concept in the biodiversity-ecosystem functioning framework to a hyperdiverse community, revealing how trophic niche partitioning can mediate biomass production across hundreds of coexisting species.

### Implications

Our findings confirm limiting similarity as a key mechanism of coexistence in one of the world’s most diverse ecosystems. The ubiquity of trophic niche partitioning across our Polynesian food web indicates that we have vastly underestimated trophic uniqueness in reef fishes, as well as overarching food web complexity on coral reefs. In addition, we show a clear, positive relationship among biodiversity, trophic partitioning, and standing biomass, demonstrating how niche-based processes can drive the relationship between biodiversity and ecosystem functioning in hyperdiverse communities when resolved with sufficiently detailed data.

Climate change poses a serious threat to coral reefs worldwide (*52*). With increasing rates of climate-induced coral bleaching, reef degradation threatens the ecosystem services that reefs provide to humans, such as food production and coastal protection, but the repercussions of coral bleaching on coexistence, community assembly, and ecosystem functioning in a coral reef ecosystem have remained unclear (*53*). Defining trophic relationships among species is essential to understand the potential of the system to buffer external perturbations and maintain the flow of energy and nutrients through coral reef communities.

Trophic specialists are highly vulnerable to global change, and the loss of a few fish consumers can have major consequences for the stability and resilience of coral reef food webs (*39*), resulting in altered energetic pathways and ecosystem collapse in previously robust networks (*54*). While global extinctions of fish species are rare, local extinctions of trophic specialists are widely documented, and communities often shift toward ecological generalization (*55*). In these scenarios, highly diverse communities will inevitably lose valuable contributors to standing stock biomass. In turn, as species richness declines, reef fish assemblages may disproportionately favor trophic generalists with high abundances and higher population resilience (*56*). Accordingly, we find that standing stock biomass is boosted in low diversity communities dominated by trophic generalists (i.e., the least amount of trophic niche segregation). The long-term consequences of repeated local extinctions, however, have yet to be assessed in historically high diversity communities (*57*).

Extreme trophic partitioning on coral reefs has implications for other hyperdiverse communities, such as tropical rainforests and freshwater systems. The tropics host over 75% of species on Earth, yet they are subjected to multiple stressors that jeopardize their biodiversity, such as deforestation, overfishing, and global change (*58*). Assuming that high biodiversity gives rise to extreme trophic partitioning in systems beyond coral reefs, they may be even more vulnerable than expected. In the ocean, it is estimated that over 90% of life remains undescribed (*14*), and marine biomes are undergoing rapid species composition changes (*59*). As we risk losing species that are not even known to science, we are also in peril of severing unique trophic connections that contribute to food web stability and sustain ecosystem services to humankind (*60*).

## Acknowledgements

We thank Titouan Roncin, Gabrielle Martineau, Kailey Bissell, Calvin Quigley, Tommy Norin, Jérémy Carlot, and Pierrick Harnay for assistance with field collections and fish dissections. We thank the staff of CRIOBE, Mo’orea for field support. We thank Anne Haguenauer for assistance with DNA extractions, and Jason Vii and Juan Pablo Lozano Peña for support with stable isotope sample preparation. We thank Lauric Thiault for providing satellite images and mapping files. Douglas Rasher provided helpful comments that improved the manuscript. All collections were performed under the guidelines and authority of the Direction de l’Environement de Polynésie Française (DIREN) under permit number 681/MCE/ENV.

## Funding

BNP Paribas Foundation (Reef Services Project; VP)

Agence Nationale de la Recherche (ANR; REEFLUX Project; ANR-17-CE32-0006; VP)

Make Our Planet Great Again Postdoctoral Grant (mopga-pdf-0000000144; JMC)

The University of Texas at Austin Marine Science Institute (JMC)

## Author contributions

Conceptualization: JMC, VP, CPM Methodology: JMC, VP

Investigation: JMC, NMDS, SJB, SD, VP, CPM, AM, FM, MZ

Data curation: JMC, VP, SJB, NMDS

Formal analysis: VP, JMC, SJB, GS

Visualization: JMC, VP, SJB, GS \

Funding acquisition: VP, JMC

Supervision: JMC, VP

Writing – original draft: JMC, VP

Writing – review & editing: all authors

## Data and materials availability

All raw sequence data generated in this study are deposited in the NCBI Sequence Read Archive (SRA) under BioProject PRJNA1522544. Individual BioSample and SRA run accessions for all 5,097 sequencing libraries are provided in Data S1. All other data and code will be made available on Dryad upon acceptance.

## Supplementary Materials

Materials and Methods

Figs. S1 to S6

Tables S1 to S5

References 1-41

Data S1

