## Supplementary Materials for "Ubiquitous trophic niche partitioning drives species coexistence and biomass on coral reefs"

**The PDF file includes:**

Materials and Methods
Figs. S1 to S6
Tables S1 to S5
References 1-41

**Other Supplementary Materials for this manuscript include the following:**

Data S1

### Materials and Methods

Field sampling

We conducted fish collections around Mo'orea, French Polynesia over five field seasons spanning three years: October 2017, March 2018, June-July 2018, October-November 2018, and May-June 2019. In total, we collected and identified 2,060 individuals across 261 species and 48 families (Table S1). We collected fishes from 132 sites across three coral reef habitats: the sheltered lagoon (0.5-10 m), exposed reef slope (5-40 m), and reef passes (5-20 m) that connect the lagoon and slope (Fig. 1). Thus, all fishes were collected between 0.5-40 m, and the collections were distributed across all reef habitats and seasons (wet and dry seasons). All cryptobenthic fishes were collected between 9:00 and 15:00. Mobile fishes were collected between 9:45 and 14:30; however, nocturnal fishes often have empty guts by mid-morning, so we also collected a subset of nocturnal mobile fishes (families: Holocentridae, Cirrhitidae, Serranidae) between 6:30 and 8:30. Following (*1*), we collected all cryptobenthic fishes with a 5:1 clove oil:ethanol solution (clove bud oil: Jedwards International, Inc., Braintree, MA, USA) via SCUBA. We collected all mobile fishes by spearfishing via freediving and SCUBA. Once brought to the surface, all fishes were immediately placed on ice. Fishes were kept on ice until dissection or preservation in the lab at the Centre de Recherches Insulaires et Observatoire de l'Environnement (CRIOBE) in Mo'orea.

Laboratory processing

In the wet lab, we measured (standard and total length), weighed, and photographed all fishes. We measured cryptobenthic fishes with digital calipers to the nearest 0.1 mm, and we took weights to the nearest 0.001 g. We photographed cryptobenthic fishes against a black or white background in a small photo tank using a Nikon D300 DSLR camera with an AF-S Micro Nikkor 60 mm macro lens (f/2.8G ED; Nikon Inc., Melville, NY, USA). We measured mobile fishes to the nearest 0.1 cm, and we weighed them to the nearest 0.1 g. All fishes were identified to species, except for two cryptobenthic fish species (the goby *Callogobius* sp. and the triplefin *Enneapterygius* sp.), which are currently taxonomically undescribed but were reliably identified based on distinct morphological features. For all other species, if we had uncertainty regarding species-level identification, they were excluded from the analyses.

We took stable isotope samples from a subset of cryptobenthic and mobile fishes. For cryptobenthic fishes, to obtain enough material for stable isotope analysis, we sampled the entire caudal region, from the anus to the caudal fin, while carefully avoiding the gastrointestinal tract. We froze the posterior half of the cryptobenthic fishes for later preparation for stable isotope analysis. We preserved the anterior half of the cryptobenthic fishes in 95% ethanol and stored the samples at -20° C for gut content sampling. We proceeded with mobile fish dissections for stable isotope and gut content sampling immediately after collection. To sample white muscle tissue from mobile fishes, we removed all scales between the caudal and dorsal fins, then we used a scalpel to extract an approximately 1 $cm^3$ section of white muscle tissue.

For gut content dissections, we used a solution of 10% bleach and sterile Milli-Q water to sterilize all surfaces and tools prior to each dissection. We removed the entire gastrointestinal tract from each fish. For mobile fishes, we extracted and homogenized all prey items from the foregut using a mortar and pestle, and then we used up to 0.3 g of the homogenized sample for DNA extraction. If the foregut was empty, we extracted more digested remnants of prey items from the hindgut and noted this caveat. We did not use the individual for gut content analysis if the entire gastrointestinal tract was empty. For cryptobenthic fishes (fishes < 5 cm; (*2*)), we performed dissections with a Zeiss V20 SteREO dissecting microscope, and the entire gastrointestinal tract was used for DNA extraction.

### Fish gut content DNA metabarcoding

#### *DNA extractions*

DNA extractions were conducted within two hours after dissection using Qiagen DNeasy PowerSoil DNA Isolation Kits (Qiagen, Hilden, Germany) at the CRIOBE in Mo'orea. Immediately after dissection, the entire alimentary tract (cryptobenthic fishes) or the homogenized gut content samples (mobile fishes) were placed in the digestion buffer in PowerBead Tubes. Before following the manufacturer's protocol, we added two 10-minute heating incubation steps to facilitate digestion, one at 65° C and another at 95° C, and then we vortexed the samples at 1,400 rpm for 10 minutes. We stored DNA extracts at -80° C and shipped the samples on dry ice to Jonah Ventures (Boulder, Colorado, USA) for amplicon-based DNA metabarcoding library preparation, sequencing, and bioinformatics.

#### *Primers*

We targeted portions of the small subunit 18S rRNA (18S), mitochondrial cytochrome *c* oxidase subunit I (COI), and chloroplast 23S rRNA (23S) gene regions. We amplified a portion of the V4 region of 18S using the V4_18SF and V4_18SR primers (*3*), the COI gene region using the mICOIintF and jgHCO2190 primers (*4*, *5*), and the 23S gene region using the p23SrV_f1 and DIam23Sr1 primers (*6*) (Table S2). The universal 18S marker served as an umbrella marker across all eukaryotic taxa, the universal COI marker provided high-resolution taxonomic information across most metazoans, and the 23S marker provided high-resolution taxonomic information across most algae. For 18S, COI, and 23S primers, forward and reverse primers contained a 5' adaptor sequences to facilitate indexing and sequencing.

#### *First PCR amplification and cleanup*

For library preparation across all markers, we ran the first PCR reactions with a volume of 25 µl following Promega PCR Master Mix specifications (Promega, Madison, WI, USA): 12.5 µl Master Mix, 0.5 µM forward and reverse primers, 1.0 µl gDNA, and 10.5 µl DNase/RNase-free water. For 18S, thermocycling followed a two-step PCR amplification. The first step included an initial denaturation at 98° C for 30 seconds, then 10 cycles of 10 seconds at 98° C and 15 seconds at 72° C. The second step included 15 cycles of 10 seconds at 98° C, 30 seconds at 62° C, and 15 seconds at 72° C, followed by a final elongation at 72° C for 7 minutes. For COI, PCR amplification included an initial denaturation at 94° C for 2 minutes, followed by 45 cycles of 15 seconds at 94° C, 30 seconds at 50° C, and 1 minute at 72° C, followed by a final elongation at 72° C for 10 minutes. For 23S, PCR amplification included the following conditions: initial denaturation at 94° C for 3 minutes, followed by 40 cycles of 30 seconds at 94° C, 45 seconds at 55° C , and 1 minute at 72° C, followed by a final elongation at 72° C for 10 minutes. Each PCR reaction was visually inspected using 5 µl of PCR product on a 2% agarose gel to determine amplicon size and ensure PCR success. We cleaned PCR amplicons by incubating amplicons with Exo1/SAP for 30 minutes at 37° C, followed by inactivation at 95° C for 5 minutes and storage at -20° C.

*Second PCR amplification, normalization, and pooling*
All remaining library preparation protocols apply across the 18S, COI, and 23S markers. We performed a second PCR amplification to give each sample a unique 12-nucleotide index sequence. This amplification followed Promega Master Mix specifications and included Promega Master Mix, 0.5 µM of each primer, and 2 µl of template of cleaned, amplified DNA from the first PCR reaction. PCR amplification included an initial denaturation of 95° C for 3 minutes, followed by 8 cycles of 95° C for 30 seconds, 55° C for 30 seconds, and 72° C for 30 seconds. Each PCR reaction was visually inspected using 5 µl of indexing PCR product input on a 2% agarose gel to ensure PCR success. We cleaned and normalized 25 µl of each indexed amplicon using the SequalPrep Normalization Kit (Life Technologies, Carlsbad, CA, USA) following the manufacturer's protocols. To pool samples, we combined 5 µl of each normalized sample.

*Sequencing*
Pooled libraries were sequenced through the Genohub service provider (Austin, TX, USA). Prior to sequencing, they performed quality control measures, including bead cleaning with AmpPure XP beads (Beckman Coulter, Brea, CA, USA) to remove non-specific < 200 bp amplicons, quantification with a Qubit 4 Fluorometer (Invitrogen, Carlsbad, CA, USA), and amplicon average size analysis with a TapeStation 4200 (Agilent, Santa Clara, CA, USA). Sequencing was performed on an Illumina HiSeq 2500 using the HiSeq Rapid SBS Kit v2 with 500 cycles (Illumina, San Diego, CA, USA). Raw sequence files were deposited in the National Center for Biotechnology Information (NCBI) Sequence Read Archive (SRA) under BioProject PRJNA1522544. A summary of samples, markers, and accessions for all sequencing libraries is provided in Data S1. Of the 5,097 sequencing libraries generated in this study, 63 libraries returned zero reads and were excluded from the submission, yielding 5,034 libraries with sequence data. These exclusions included 39 fish samples: 31 had partial marker dropout but were retained, while eight had no data across any marker and were excluded from all downstream analyses.

<u>Sequence bioinformatics</u>
For 18S, COI, and 23S markers, we demultiplexed raw sequence data using pheniqs v2.1.0 (*7*), using strict matching of sample barcode indices with no errors. We then used Cutadapt v3.4 (*8*) to remove gene primers from forward and reverse reads. Read pairs were discarded when one or both primers were not found at the expected 5' location with an error rate of < 0.15. Read pairs were merged using vsearch v2.15.2 (*9*). We discarded sequences with a length of less than 330 bp or greater than 450 bp for 18S, less than 298 bp or greater than 328 bp for COI, and less than 300 bp or greater than 380 bp for 23S. For all markers, we discarded sequences with an error rate of greater than 0.5 bp (*10*). Across all markers, reads for each sample were clustered using the unoise3 denoising algorithm in VSEARCH (*11*), using an alpha of 5 and discarding unique raw sequences observed less than 8 times. We then compiled counts of amplicon sequence variants (ASVs) and putative chimeras were removed using the *uchime3* algorithm through VSEARCH. For each ASV, we assigned consensus taxonomy using custom best-hits algorithm. For the 18S marker, we used the SILVA v138.1 (*12*) reference database, and for COI and 23S, we used publicly accessible GenBank (*13*) and Jonah Ventures voucher sequence records as reference databases. To search reference databases for each marker dataset, we used an exhaustive semi-

global pairwise alignment with VSEARCH. We further employed a custom, query-centric approach to quantify match quality. Here, the percent match ignores terminal gaps in the target sequence but not in the query sequence. Finally, we generated a consensus taxonomy using either all 100% matching reference sequences or including all reference sequences within 1% of the top match. For any taxonomic level, we accepted reference taxonomy with a greater than 90% agreement across the top hits.

For the COI sequences, to maximize ASVs with taxonomic assignments, we re-assigned taxonomy using a local reference database curated from the Mo'orea BIOCODE Project, a comprehensive barcode library that includes most metazoans on and around Mo'orea (*14*, *15*). We accepted genus and phylum-level taxonomy at 97% and ≥ 80% sequence similarity, respectively. Between the GenBank and BIOCODE taxonomic assignments, we selected final taxonomic assignments based on highest sequence similarity. When sequence similarity was equal, we selected the taxonomic assignment with higher taxonomic resolution.

For the COI, 18S, and 23S datasets, we standardized all taxonomic assignments using the R package *taxize* (*16*), which allowed us to develop automated scripts that fill in missing taxonomic ranks (genus, family, class, order, and phylum) by accessing two public databases with extensive taxonomic catalogues: NCBI and the World Register of Marine Species (WoRMS). We removed all singleton ASVs before proceeding with downstream data analysis.

For the COI and 18S datasets, we took a conservative approach to removing "self-hits" of fish species from the data. We removed all ASVs with taxonomic assignments that matched the species, genus, or family of the fish host. Consequently, our dataset does not capture potential cannibalism or the consumption of fishes in the same genus or family as the fish host. The 23S dataset did not require the removal of "self-hits" because it is specific to marine algae and does not amplify marine fishes.

Stable isotope analysis

For the cryptobenthic fish samples, we first decalcified them to remove bones. We crushed the cryptobenthic fishes with a bead mill, then we added 1% hydrochloric acid to each sample. After 24h, we centrifuged the samples, removed the supernatant, and rinsed the pellet three times with Milli-Q water. We repeated this decalcification process at least three times for each sample, following established protocols for cryptobenthic fishes (*17*). For mobile fishes, we stored white muscle samples at -20° C for at least 24h. Then, we freeze-dried all samples for at least 24h. After lyophilization, all samples were transported to the CRIOBE in Perpignan, France, where we homogenized them into a fine powder. We shipped homogenized samples to the Cornell Isotope Laboratory (COIL) for bulk stable isotope analysis of carbon and nitrogen. All analyses were performed on a Thermo Delta V isotope ratio mass spectrophotometer interfaced to a NC2500 elemental analyzer. Elemental percentages are calculated based on sample weight. COIL routinely calibrates their in-house standards against international reference materials from the International Atomic Energy Association. They ensure precision across sample runs and over time by analyzing an in-house standard after every 10 samples, and they calibrate their instrument across a gradient of amplitude intensities using a Methionine standard. They use two in-house standards to perform isotope corrections with a two-point normalization of all $\delta^{15}N$ and $\delta^{13}C$ data.

Reef fish community composition

Integrating historical fish community data allows us to contextualize niche segregation across entire reef fish assemblages. As part of the CRIOBE long-term monitoring program within the French Service National d'Observation (SNO) CORAIL, fish and benthic communities have been surveyed around Mo'orea, French Polynesia annually between January to June since 2004. For our analyses, we obtained fish community data from 19 years: 2005 to 2023. The monitoring involves thirteen sites, including eight marine protected areas (MPAs) and five sites outside of MPAs. For the fish surveys, three distinct reef habitats are surveyed: the fringing reef, barrier reef, and outer slope (at 10 m depth). At each habitat, standardized visual surveys of the mobile and site-attached fishes, including species identity and size, are conducted along three permanent 25x2 m belt transects. Each transect is separated by 25 m at each site. With three replicate transects per site and habitat, a total of 117 transects are run per sampling period.

Data analysis

We conducted statistical analyses using Python version 3.14.6 (*18*) and R version 4.4.2 (*19*).

*Molecular and isotopic data preparation*

To combine the COI, 18S, and 23S metabarcoding datasets, we adopted a conservative approach to estimate niche segregation. We converted sequence reads to presence-absence data (zeros and ones) to minimize biases across three molecular markers, which have distinct taxonomic targets, different amplification efficiency, and variable sequencing depth. We only retained ASVs with taxonomic assignments to at least the level of phylum. We then merged the three molecular datasets based on lowest taxonomic rank, including multiple hierarchical taxonomic levels from phylum to species, into one data frame, creating a single presence-absence dataset. This multi-level approach allowed us to test whether patterns of niche segregation are robust across taxonomic resolution or emerge only at specific levels of resource classification. Our approach may overlook prey items due to primer limitations or imperfect reference databases. However, it is consistent across loci and minimizes amplification and sequencing biases that can generate disparities in sequence reads across ASVs and ultimately over or under represent particular taxonomic groups. For the isotope dataset, isotopic niches were described in continuous two-dimensional trophic space defined by $\delta^{13}C$ and $\delta^{15}N$ values measured at the individual level.

DNA metabarcoding and stable isotope analysis characterize trophic niches at different temporal and ecological scales, so we applied different minimum sample size thresholds for each dataset. DNA metabarcoding provides a taxonomically resolved snapshot of recently consumed prey, so every sampled individual provides information on the trophic niche of a species (*20*). In contrast, stable isotopes integrate assimilated diet over longer time windows (e.g., months), so isotopic niches are population-level properties estimated from the dispersion of multiple individuals in $\delta^{13}C$–$\delta^{15}N$ space, and trophic niche calculations are sensitive to small sample sizes even after small sample correction (*21*). Accordingly, species were retained if represented by at least two individuals for metabarcoding and at least five individuals for stable isotopes.

*Trophic network*

Using the metabarcoding dataset, we visualized the bipartite network linking fish species to their prey items using a hierarchical edge-bundling layout implemented in Python with Matplotlib

(*22*), igraph (*23*), and scikit-learn (*24*). Following (*25*), fish species were grouped into eight trophic groups and arranged around an outer ring. Prey nodes were clustered into groups of taxonomically or ecologically similar prey using agglomerative clustering (scikit-learn) on the biadjacency matrix, and positioned in the interior. Each prey cluster was attracted toward the groups in which it occurred most frequently, calculated as the proportion of fish hosts within each group in which the prey item was detected, consistent with the presence-absence structure of the dataset. To reduce visual noise from trophically rare or incoherent links (likely reflecting secondary consumption; e.g., algae in the gut of a carnivore), we applied a filter to remove all links between a food macro-category and trophic group when that category represented less than $\theta = 0.03$ (i.e., 3%) of the total diet of that group. Edges were drawn as Bézier curves routed through shared intermediate control points between communities, producing the bundling effect that highlights dominant trophic pathways. For visualization, prey nodes were colored by six prey categories: algae, macroinvertebrates, microbes, microinvertebrates, sessile invertebrates, vertebrates, which were assigned based on prey taxonomy (Table S3).

*Intraspecific and interspecific trophic niche segregation*

We first examined whether niche overlap primarily emerges within species or among species, a central question of individual specialization theory (*26*, *27*). Under this framework, population-level niches arise from the aggregation of heterogenous individual niches, and interspecific differentiation is expected to dominate only when within-species niche variation is limited. To test this, we quantified niche segregation by explicitly analyzing within-species (intraspecific) and between-species (interspecific) variation.

For the metabarcoding and isotope datasets, intraspecific niche segregation was quantified as niche variation among individuals belonging to the same species. For the metabarcoding data, we calculated the average pairwise dissimilarity in prey composition among conspecific individuals with two complementary overlap-based indices: (i) a Simpson-based turnover ($\beta$sim; (*28*)) and (ii) the inverse of Pianka's index, calculated as one minus Pianka overlap, adapted for presence–absence data (*29*). Pianka's index is widely used to quantify trophic niche overlap, but it is sensitive to niche breadth and may inflate overlap values when diets are broad (*30*). In contrast, $\beta$sim partitions beta diversity into turnover and nestedness components and specifically quantifies pure turnover, making it independent of differences in niche breadth or prey richness (*28*). Interspecific niche segregation was quantified using the same overlap-based metrics to ensure direct comparability with intraspecific estimates. We detected high correlation between the Simpson and Pianka metrics (Pearson correlation coefficient: $r = 0.913$), so we only report Pianka-based metrics.

Overlap-based indices (e.g., Simpson, Pianka) are designed for discrete resource-use data (e.g., itemized gut contents) and are therefore not directly applicable to trophic positions in continuous isotopic space. Thus, for the isotope data, intraspecific niche segregation was quantified as the mean pairwise Euclidean distance among conspecific individuals in $\delta^{13}C$–$\delta^{15}N$ space, calculated across all individual pairs within each species. This metric captures the degree of trophic dispersion among individuals without requiring an explicit definition of niche overlap. In contrast, interspecific isotopic niche segregation was quantified as the Euclidean distance between species centroids in the isotopic space, where each centroid was defined by the mean $\delta^{13}C$ and $\delta^{15}N$ values across individuals of a species. Using centroid-to-centroid distances provides a parsimonious measure of trophic niche segregation among species while reducing sensitivity to within-species isotopic variance.

*Null modelling of trophic niche segregation*

To assess whether observed intraspecific segregation exceeded random expectations, we evaluated both metabarcoding and isotopic metrics against null models. Null models were designed to remove species-specific structure while preserving the distribution of trophic information at the community level. For the metabarcoding data, null expectations were generated by randomly pairing individuals drawn from the entire consumer pool, irrespective of species identity. For each random pair, dietary niche segregation was computed using the same Pianka-based metrics applied to conspecific individuals. This null model preserves the global frequency of prey occurrences and the distribution of individual dietary richness but randomizes the species identity of individuals. As such, it tests whether individuals belonging to the same species are more (or less) partitioned in their diets than expected by chance given the overall heterogeneity of individual diets in the community. For the isotope data, intraspecific null expectations were constructed for each species by repeatedly sampling the same number of individuals from the isotope dataset, ignoring species identity. Mean pairwise Euclidean distances were then calculated for each resampled group. This null model preserves the empirical isotopic distribution of the community while removing species-level clustering in isotopic space, allowing us to test whether observed within-species isotopic dispersion exceeds that expected by chance (i.e., randomly grouping individuals).

Interspecific null models were constructed to test whether observed trophic niche segregation among species exceeds expectations by chance. For the metabarcoding data, null expectations were generated using a fixed-marginal randomization of the prey presence–absence matrix based on the Curveball algorithm (*31*). This procedure preserves row sums (prey occurrence frequencies) and column sums (species dietary richness), while randomizing the specific associations between consumers and prey. As a result, the null model maintains the empirical distribution of niche breadth for consumers and prey commonness across the community, while removing non-random patterns of dietary niche partitioning. Pairwise interspecific segregation was calculated for each randomized matrix using the same overlap-based metrics used for the observed data. For the isotope data, interspecific null expectations were generated by permuting species identities among individuals prior to centroid estimation. We calculated species centroids for each permutation, then computed Euclidean distances between centroids. This null model preserves the isotopic structure of the species pool and empirical distribution of individual isotopic values, while removing species-level aggregation in isotopic space. It therefore tests whether observed trophic segregation among species exceeds that expected by chance (i.e., randomly assigning individuals to species).

Null model selection reflects fundamental differences in the data structure of the metabarcoding and isotope datasets, as well as the definition of niche segregation across organizational levels. For the metabarcoding data, interspecific segregation is evaluated against null models that preserve totals in discrete species–resource matrices, ensuring that differences in dietary breadth and prey availability do not drive niche segregation. For isotopic data, niches are defined in continuous space, so null models preserve the empirical variance structure while removing species-specific clustering. Together, these null models allow a consistent assessment of whether intra and interspecific niche segregation reflect non-random trophic structuring rather than artefacts of sampling, niche breadth, or marginal constraints.

*Community-level trophic niche segregation*
To assess whether local fish communities around Mo'orea assemble with non-random trophic structure, we quantified trophic niche segregation among co-occurring species from the CRIOBE long-term monitoring data collected from 2005-2023 (see Methods: Reef fish community composition). From this dataset, we derived species-specific biomass estimates, local community composition, and reef habitat specific (i.e. fringing reef, barrier reef, and outer slope) species pools. For both the metabarcoding and isotope data, we calculated average trophic niche segregation among species within a community (i.e., a site) and compared observed patterns to null expectations derived from random assemblages drawn from the regional species pool. For each transect, we only considered species observed in underwater visual censuses for which trophic niche information was available. Under our previously defined thresholds of removing species with less than two individuals from the metabarcoding dataset and five individuals from the isotopic dataset (see *Molecular and isotopic data preparation*), retained species accounted for most of the total fish biomass per transect in the metabarcoding dataset (86% ± 15% SD) and a smaller fraction in the isotope dataset (42% ± 22% SD; Fig. S2).

Trophic niche segregation was quantified for all pairs of co-occurring species using two complementary overlap-based metrics. For dietary metabarcoding data, individual diets were aggregated at the species level, then pairwise niche segregation was computed using the same Simpson-based turnover (βsim) and the Pianka-based index as previously described. For the isotope data, there are no existing methods to compute these two niche segregation indices. We therefore developed custom algorithms to compute them from Bayesian standard ellipses generated in isotopic $\delta^{13}C$–$\delta^{15}N$ space using the R package *nicheROVER* (*32*). Precisely, for each species pair, a two-dimensional sampling domain encompassing both ellipses was defined based on their centroids and covariance structure. A large number of random points (500 per square unit of isotopic space) were uniformly drawn in isotopic space, and points were classified as belonging to one or both ellipses based on their Mahalanobis distance relative to the $\chi^2$ threshold associated with α (set to 0.05, corresponding to 95% ellipses). We then estimated ellipse areas (A and B) and their shared area (C) from the proportion of sampled points falling within each region. Once these area estimates were obtained, we evaluated pairwise niche segregation using two indices: (i) Simpson-based turnover (βsim) was calculated as $\min(A - C, B - C)/(C + \min(A - C, B - C))$, which isolates the turnover component of isotopic differentiation and is independent of differences in niche breadth, and (ii) Pianka segregation was calculated as $1 - C/\sqrt{(AB)}$ (Fig. S4). These metrics provide isotopic analogues of the overlap-based indices used for the metabarcoding data, allowing direct comparison across the two datasets.

At the community level, we summarized pairwise niche segregation in two ways. First, we calculated mean niche segregation across all species pairs, treating each pair as contributing equally to community trophic structure. Second, we calculated biomass-weighted mean niche segregation, where each species pair was weighted by their relative biomass within each transect. The biomass-weighted metric reflects an ecologically realistic scenario where trophic structure and competition for resources is shaped not only by consumer presence, but also by the relative effects on resource consumption. Accordingly, niche segregation among dominant species is expected to have a disproportionate influence on resource partitioning, interaction strength, and energy flow within the community, whereas niche segregation involving rare species contributes less to overall community-level trophic organization. By contrasting unweighted and weighted metrics, we explicitly test whether interactions between trophic niche partitioning, coexistence, and biomass are driven by dominant consumers or emerge equally across species pairs irrespective of abundance.

To evaluate whether observed community-level niche segregation differed from random chance, we implemented a transect-specific null model that randomizes species co-occurrence while preserving key ecological constraints. For each transect, we generated null communities by randomly drawing the same number of species from a habitat-specific regional pool, defined as all species observed in the same reef zone (i.e., outer reef, barrier reef, and fringing reef) and for which trophic niche data were available. Importantly, species-level trophic niches were fixed during randomization. For each null community, observed biomass values were reassigned to randomly selected species, thereby preserving the observed biomass distribution within the transect while randomizing over species identities, trophic niches, and local co-occurrence. Community-level trophic niche segregation was then calculated for each null assemblage using the same unweighted and biomass-weighted metrics as used for the observed data.

Observed community-wide niche segregation values were standardized as z-scores relative to the null distributions (i.e., mean(observed) – mean(null) / SD(null)). This allowed us to test whether local communities exhibit higher or lower trophic segregation than expected under random assembly from the Mo'orea species pool. We detected a high correlation between z-scores from the Simpson and Pianka metrics, suggesting that differences in niche breadth between species pairs do not drive the observed patterns (Fig. S5).

*Trophic niche segregation and realized total biomass*

We expected species richness and trophic niche segregation to jointly shape community biomass due to the interactive effects of complementarity and biodiversity on ecosystem functioning (*33*, *34*). Species richness can increase standing biomass through niche complementarity and more complete resource use (*35*). However, the potential for complementarity-drive effects may also be shaped by how many species locally co-occur: in species-rich assemblages, competition for shared resources can be stronger and niche differences may become more consequential, potentially amplifying the positive association between segregation and biomass (*36*, *37*). Conversely, if species in diverse communities force trophic overlap, niche segregation may contribute little to biomass gains at high diversity, leading to weaker or saturating effects (*38*).

To examine this, we analyzed the interactive effects of species richness and community-level niche space occupation on community-level biomass. Specifically, we built Bayesian models to predict transect-level fish standing biomass as a function of richness, niche segregation, and their interaction, using reef habitat as a covariate. Given the high correlation between the Simpson and Pianka metrics (Fig. S5), we only used Pianka-based indices. We built models using unweighted and biomass-weighted metrics of niche segregation to account for the possibility that trophic differentiation among dominant, high-biomass species disproportionately influences community-level biomass. Biomass and richness were log-transformed and standardized prior to modelling, and we used the previously computed z-scores for niche segregation metrics from the community-level null models.

We evaluated potential multicollinearity at the model level using the condition index, which was calculated from linear models including richness, segregation, their interaction, and reef habitat as a covariate. The condition index was chosen over the variance inflation factor because the metric is appropriate in the presence of interaction terms, for which variance inflation factors can become inflated even under moderate correlation. Across all datasets and segregation scenarios, condition indices remained below commonly used thresholds for severe multicollinearity (i.e., CI $< 30$), indicating that collinearity was unlikely compromising parameter estimation (Table S4). However, the Pearson correlation coefficient was high between metabarcoding-based biomass-weighted segregation and species richness ($r = -0.78$), the correlation between isotope-based biomass-weighted segregation and species richness was weaker ($r = -0.58$), and the correlation between unweighted segregation metrics and species richness was much weaker ($r < 0.3$).

Models were fitted with the R package *brms* (*39*) with a Gaussian error distribution and an identity link using an interface to Stan with the R package *cmdstanr* (*40*). The model structure follows:

$$\text{Biomass} \sim \text{Richness} \times \text{Segregation} + \text{ReefZone} \qquad (1)$$

where ReefZone was included as a categorical covariate to account for habitat-related differences in biomass. Segregation is represented by z-scores from: (i) metabarcoding biomass-weighted niche segregation, (ii) metabarcoding unweighted niche segregation, (iii) isotope biomass-weighted niche segregation, and (iv) isotope unweighted niche segregation. For each dataset, we fitted paired models with and without the richness × segregation interaction to test whether the effect of niche segregation on biomass depends on local species richness. We evaluated the support for interaction terms using approximate leave-one-out cross-validation (LOO), which compared models with and without the richness × segregation interaction within each dataset and segregation metric (Table S5).

All models were estimated using weakly informative priors to regularize parameter estimates while remaining agnostic about effect direction and magnitude. Regression coefficients (including main effects and interactions) were assigned normal priors centered at zero (mean = 0, SD = 1). Intercepts were assigned normal priors (mean = 0, SD = 1), and residual standard deviations were assigned Student-t priors with 3 degrees of freedom, mean 0, and scale 1. Models were run with 4 Markov chains, with 8,000 iterations per chain, and a warm-up of 50%. Sampling was performed using the No-U-Turn Sampler (NUTS), with an *adapt_delta* of 0.99 and a maximum tree depth of 15 to ensure stable exploration of the posterior. Convergence was assessed using R-hat statistics (all $< 1.01$), effective sample sizes (ESS), and visual inspection of trace plots. We assessed model fit with posterior predictive checks (Fig. S6).

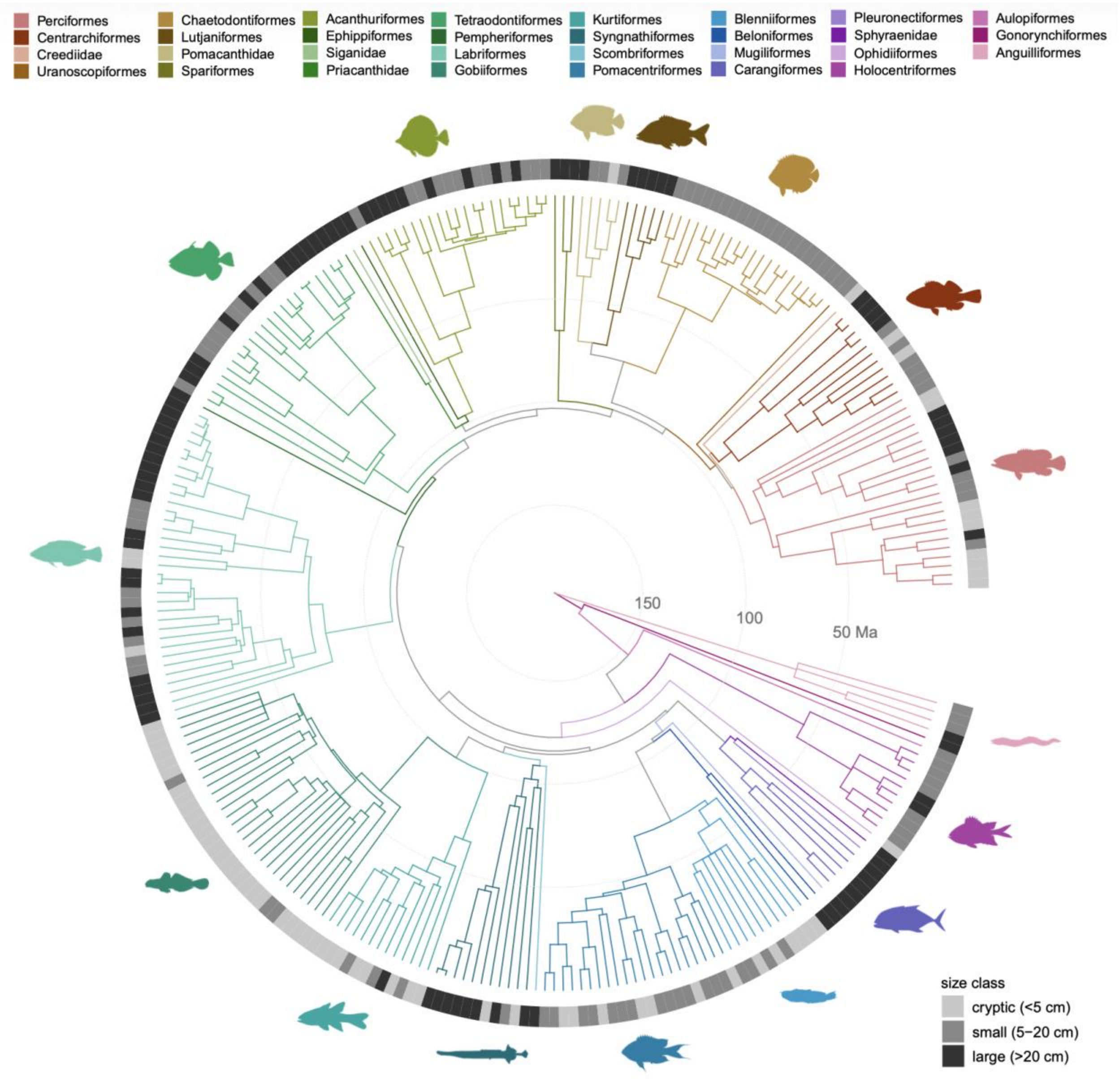


**Fig. S1.**

Body size across the phylogeny of fish species in this study. Time-calibrated phylogeny of the 261 fish species included in this study, pruned from the Fish Tree of Life chronogram (*41*). Branches are colored by order, with fish silhouettes representing each clade. The outer ring indicates body-size classes in grayscale.

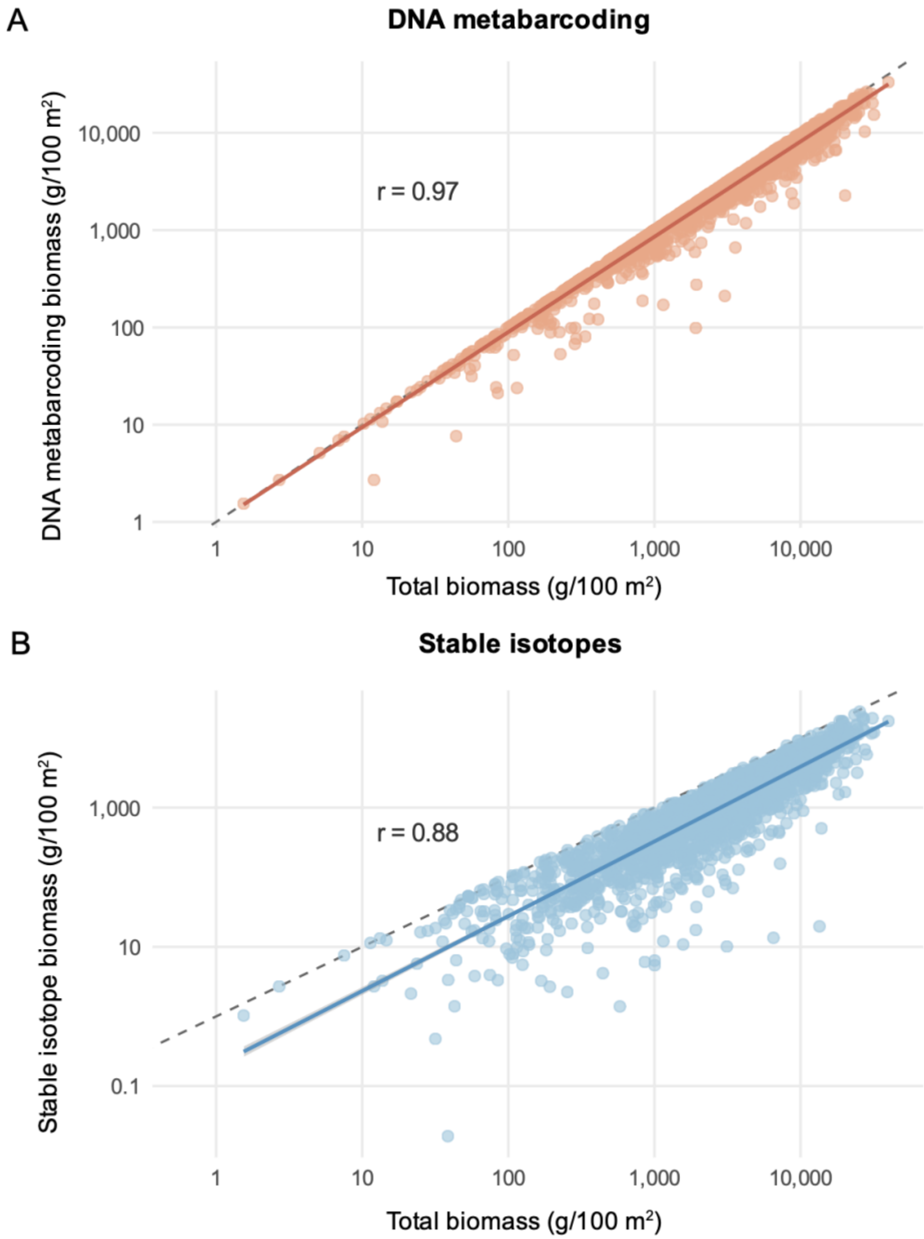


**Fig. S2.**

Biomass captured by the molecular and isotopic analyses relative to total fish biomass. Biomass included in the (**A**) DNA metabarcoding and (**B**) stable isotope trophic niche segregation analyses plotted against total fish biomass per transect from the long-term monitoring data. Each point represents fish biomass from one transect. Solid lines show fitted regressions, while dashed lines represent the 1:1 relationship expected under perfect correspondence between included and measured fish biomass. Pearson correlation coefficients (r) are shown for each panel.

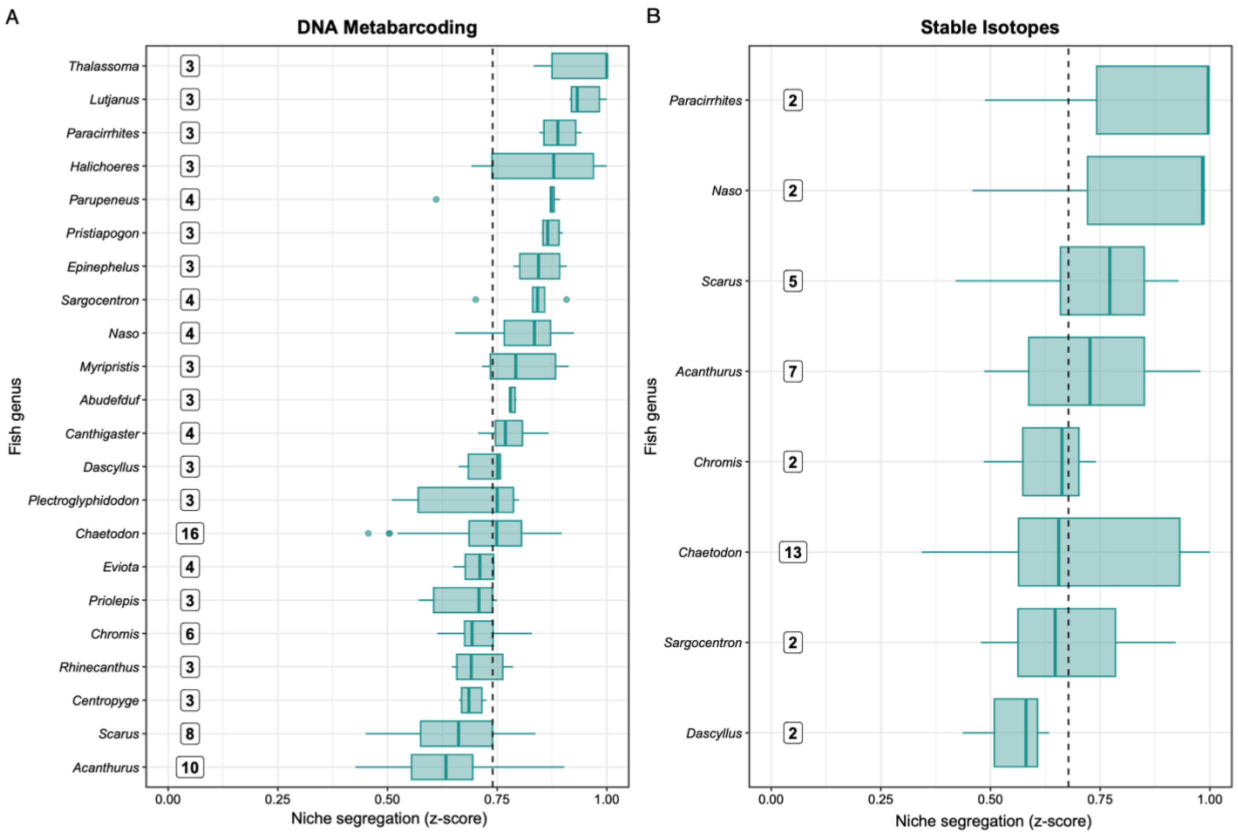


**Fig. S3.**

Trophic niche partitioning across fish species based on molecular and isotopic approaches. Percent niche segregation (z-scores) between species pairs, within fish genera, based on (**A**) Pianka overlap for molecular and (**B**) Euclidean distances for isotopic data. The numbers to the left of the boxplots indicate the total number of species sampled within that family. The boxplots represent the medians and interquartile ranges, and the whiskers represent 1.5 times the interquartile range. Points outside the boxplot whiskers represent outliers. Fish genera are ranked from highest (top) to lowest (bottom) trophic niche segregation based on median values. The dashed vertical lines represent the average niche segregation of all species across the (**A**) 22 fish genera in the molecular dataset and (**B**) and 8 fish genera in the isotopic dataset.

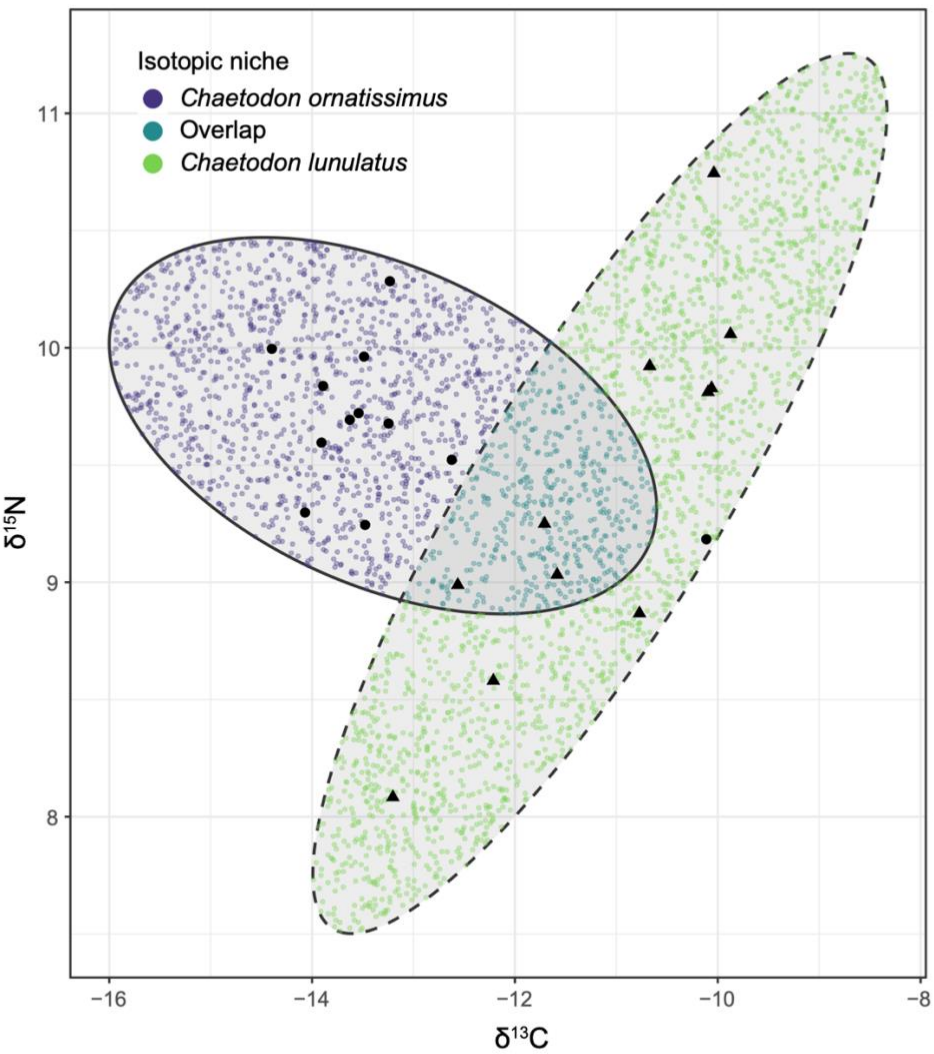


**Fig. S4.**
Isotopic niche partitioning between two butterflyfishes: *Chaetodon ornatissimus* and *Chaetodon lunulatus*. Solid and dashed lines represent the 95% isotopic niche ellipses in $\delta^{13}C$–$\delta^{15}N$ space for *C. ornatissimus* and *C. lunulatus*, respectively. Black circles and triangles represent $\delta^{13}C$ and $\delta^{15}N$ measurements of individual *C. ornatissimus* and *C. lunulatus*, respectively, while colored points show simulated values used to estimate ellipse area and overlap.

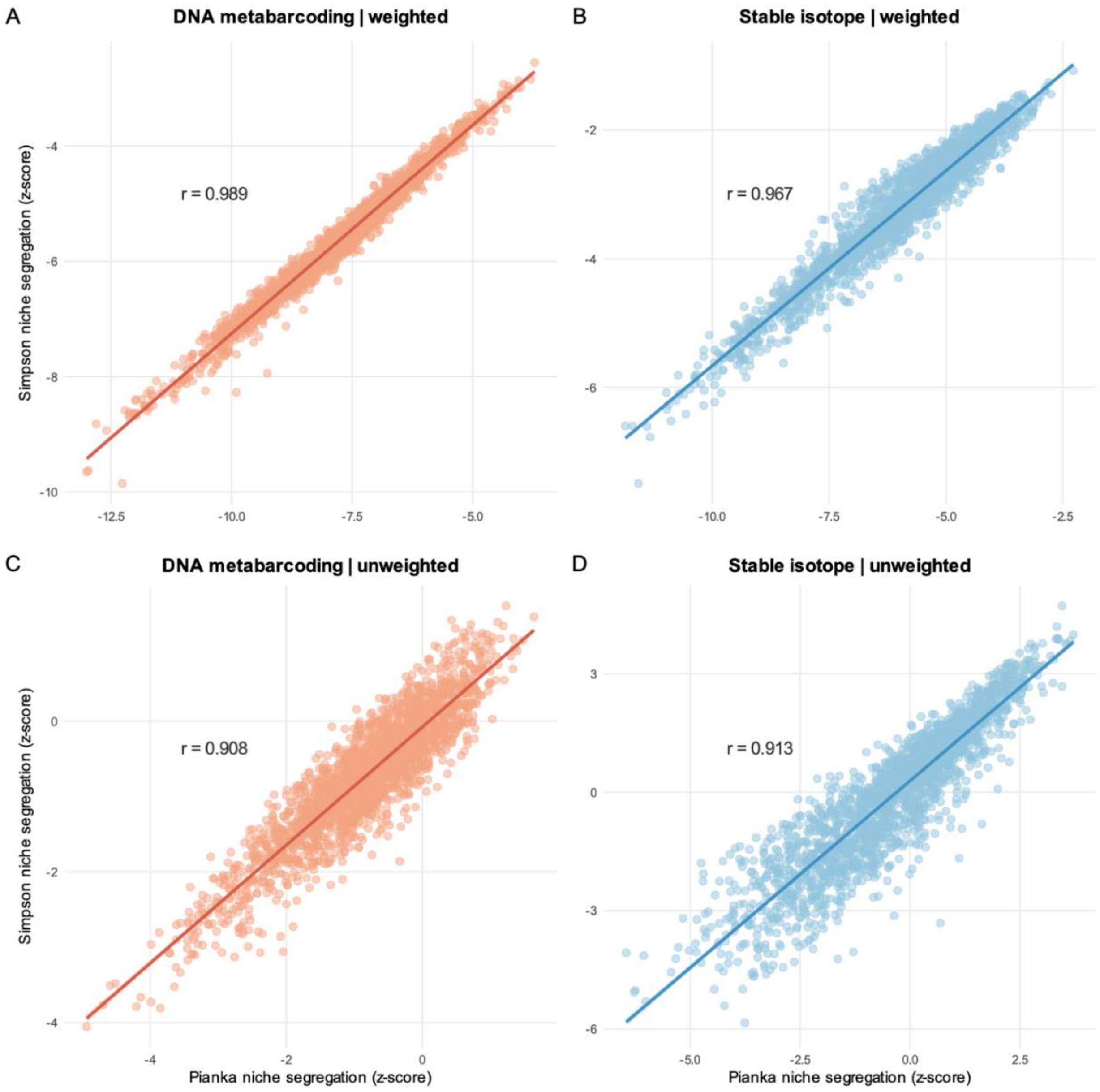


**Fig. S5.**

Correlation between two indices of pairwise niche segregation. Simpson turnover (βsim) and Pianka overlap for (**A**, **C**) DNA metabarcoding and (**B**, **D**) stable isotope data, under (**A** to **B**) biomass-weighted and (**C** to **D**) unweighted formulations. Both metrics are shown as null-model-standardized z-scores. Each point represents one pairwise species comparison, and solid lines show fitted regressions. Pearson correlation coefficients (r) are shown for each panel.

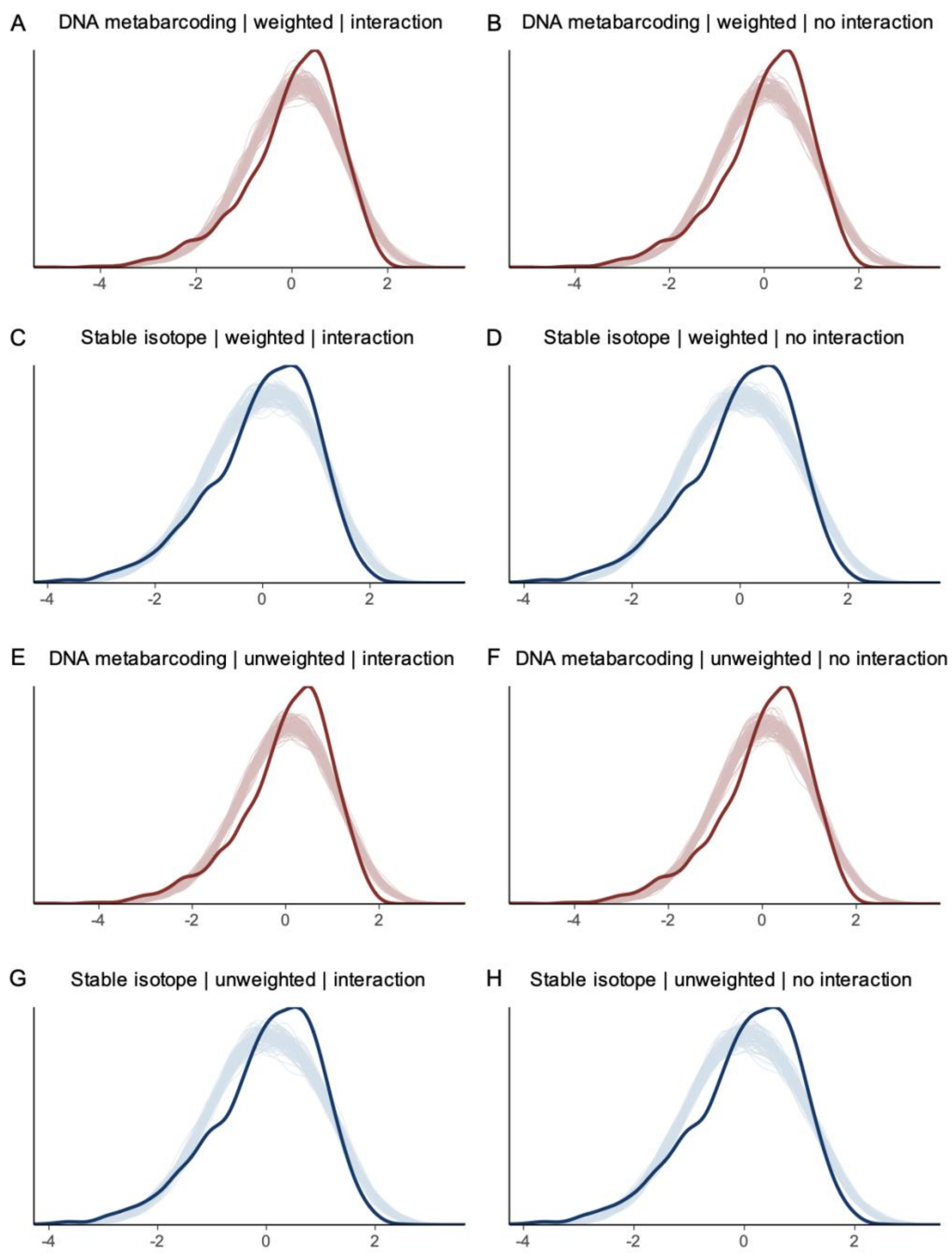


**Fig. S6.**

Posterior predictive checks for Bayesian models of niche segregation. For each model, the observed distribution of standardized transect-level fish biomass (dark red and blue lines) is compared to draws simulated from the posterior predictive distribution (light red and blue lines). Panels show all combinations of data types: (**A** to **B**, **E** to **F**) DNA metabarcoding (red), (**C** to **D**, **G** to **H**) stable isotope (blue); weighting schemes: (**A** to **D**) weighted, (**E** to **F**) unweighted; and the inclusion of an interaction term, as indicated by the panel titles. Posterior predictions closely match the empirical data, indicating no systematic lack of fit.

**Table S1.**

Sampling effort and trophic guilds of fish species included in this study. Summary of the 2,060 individuals across the 261 fish species collected, identified, and analyzed for gut content DNA metabarcoding and stable isotope analysis, including their assigned trophic guild ((*25*); HMD = herbivore, microvore, detritivore). Metabarcoding and Isotope columns indicate the number of individuals analyzed per species for each method. A total of 260 species were analyzed for gut content DNA metabarcoding, and a subset of 216 fish species were analyzed using bulk stable isotope analysis.

| Family | Species | Trophic guild | Metabarcoding | Isotope |
|---|---|---|---|---|
| Acanthuridae | *Acanthurus_achilles* | HMD | 10 | 9 |
| Acanthuridae | *Acanthurus_guttatus* | HMD | 10 | 10 |
| Acanthuridae | *Acanthurus_lineatus* | HMD | 10 | 10 |
| Acanthuridae | *Acanthurus_nigricans* | HMD | 11 | 10 |
| Acanthuridae | *Acanthurus_nigrofuscus* | HMD | 9 | 9 |
| Acanthuridae | *Acanthurus_nigros* | HMD | 5 | 5 |
| Acanthuridae | *Acanthurus_olivaceus* | HMD | 10 | 6 |
| Acanthuridae | *Acanthurus_pyroferus* | HMD | 10 | 8 |
| Acanthuridae | *Acanthurus_thompsoni* | planktivore | 5 | 4 |
| Acanthuridae | *Acanthurus_triostegus* | HMD | 12 | 2 |
| Acanthuridae | *Ctenochaetus_binotatus* | HMD | 1 | 1 |
| Acanthuridae | *Ctenochaetus_flavicauda* | HMD | 10 | 10 |
| Acanthuridae | *Ctenochaetus_hawaiiensis* | HMD | 1 | 0 |
| Acanthuridae | *Ctenochaetus_striatus* | HMD | 10 | 2 |
| Acanthuridae | *Naso_brevirostris* | HMD | 6 | 6 |
| Acanthuridae | *Naso_lituratus* | HMD | 10 | 10 |
| Acanthuridae | *Naso_unicornis* | HMD | 4 | 4 |
| Acanthuridae | *Naso_vlamingii* | HMD | 5 | 5 |
| Acanthuridae | *Zebrasoma_scopas* | HMD | 10 | 10 |
| Acanthuridae | *Zebrasoma_velifer* | HMD | 9 | 8 |
| Apogonidae | *Apogon_coccineus* | microinvertivore | 1 | 0 |
| Apogonidae | *Apogon_crassiceps* | microinvertivore | 8 | 0 |
| Apogonidae | *Apogon_susanae* | microinvertivore | 10 | 7 |
| Apogonidae | *Cheilodipterus_macrodon* | crustacivore | 2 | 2 |
| Apogonidae | *Cheilodipterus_quinquelineatus* | crustacivore | 7 | 0 |
| Apogonidae | *Fowleria_marmorata* | microinvertivore | 8 | 3 |
| Apogonidae | *Fowleria_vaiulae* | microinvertivore | 5 | 0 |
| Apogonidae | *Nectamia_savayensis* | planktivore | 6 | 2 |
| Apogonidae | *Ostorhinchus_angustatus* | microinvertivore | 10 | 9 |
| Apogonidae | *Ostorhinchus_nigrofasciatus* | crustacivore | 10 | 0 |

| | | | | |
|---|---|---|---|---|
| Apogonidae | *Pristiapogon_exostigma* | crustacivore | 8 | 11 |
| Apogonidae | *Pristiapogon_fraenatus* | microinvertivore | 5 | 2 |
| Apogonidae | *Pristiapogon_kallopterus* | crustacivore | 6 | 10 |
| Aulostomidae | *Aulostomus_chinensis* | piscivore | 11 | 10 |
| Balistidae | *Balistapus_undulatus* | macroinvertivore | 10 | 2 |
| Balistidae | *Balistoides_viridescens* | macroinvertivore | 1 | 1 |
| Balistidae | *Melichthys_niger* | HMD | 10 | 10 |
| Balistidae | *Melichthys_vidua* | HMD | 10 | 10 |
| Balistidae | *Odonus_niger* | macroinvertivore | 10 | 1 |
| Balistidae | *Pseudobalistes_flavimarginatus* | macroinvertivore | 1 | 0 |
| Balistidae | *Pseudobalistes_fuscus* | macroinvertivore | 1 | 0 |
| Balistidae | *Rhinecanthus_aculeatus* | microinvertivore | 10 | 5 |
| Balistidae | *Rhinecanthus_lunula* | microinvertivore | 3 | 3 |
| Balistidae | *Rhinecanthus_rectangulus* | microinvertivore | 10 | 7 |
| Balistidae | *Sufflamen_bursa* | macroinvertivore | 12 | 9 |
| Belonidae | *Tylosurus_crocodilus* | piscivore | 2 | 1 |
| Blenniidae | *Aspidontus_taeniatus* | planktivore | 3 | 0 |
| Blenniidae | *Blenniella_paula* | HMD | 3 | 0 |
| Blenniidae | *Cirripectes_variolosus* | HMD | 10 | 9 |
| Blenniidae | *Enchelyurus_ater* | HMD | 10 | 4 |
| Blenniidae | *Exallias_brevis* | HMD | 7 | 5 |
| Blenniidae | *Glyptoparus_delicatulus* | HMD | 16 | 10 |
| Blenniidae | *Plagiotremus_tapeinosoma* | HMD | 3 | 0 |
| Bothidae | *Bothus_mancus* | piscivore | 3 | 3 |
| Bythitidae | *Diancistrus_katrineae* | na | 1 | 0 |
| Caesionidae | *Pterocaesio_tile* | planktivore | 2 | 2 |
| Callionymidae | *Callionymus_filamentosus* | microinvertivore | 5 | 0 |
| Carangidae | *Caranx_ignobilis* | piscivore | 1 | 1 |
| Carangidae | *Caranx_melampygus* | piscivore | 5 | 3 |
| Carangidae | *Elagatis_bipinnulata* | piscivore | 1 | 0 |
| Carangidae | *Scomberoides_lysan* | piscivore | 3 | 3 |
| Carangidae | *Trachinotus_baillonii* | piscivore | 1 | 0 |
| Chaetodontidae | *Chaetodon_auriga* | corallivore | 10 | 6 |
| Chaetodontidae | *Chaetodon_bennetti* | corallivore | 3 | 3 |
| Chaetodontidae | *Chaetodon_citrinellus* | corallivore | 10 | 5 |
| Chaetodontidae | *Chaetodon_ephippium* | corallivore | 10 | 10 |
| Chaetodontidae | *Chaetodon_lunula* | microinvertivore | 10 | 10 |
| Chaetodontidae | *Chaetodon_lunulatus* | corallivore | 11 | 11 |
| Chaetodontidae | *Chaetodon_mertensii* | corallivore | 5 | 4 |

| | | | | |
|---|---|---|---|---|
| Chaetodontidae | *Chaetodon_ornatissimus* | corallivore | 19 | 12 |
| Chaetodontidae | *Chaetodon_pelewensis* | corallivore | 10 | 5 |
| Chaetodontidae | *Chaetodon_quadrimaculatus* | corallivore | 10 | 9 |
| Chaetodontidae | *Chaetodon_reticulatus* | corallivore | 20 | 12 |
| Chaetodontidae | *Chaetodon_trichrous* | planktivore | 6 | 5 |
| Chaetodontidae | *Chaetodon_trifascialis* | corallivore | 7 | 5 |
| Chaetodontidae | *Chaetodon_ulietensis* | corallivore | 8 | 8 |
| Chaetodontidae | *Chaetodon_unimaculatus* | corallivore | 10 | 9 |
| Chaetodontidae | *Chaetodon_vagabundus* | corallivore | 10 | 5 |
| Chaetodontidae | *Forcipiger_flavissimus* | microinvertivore | 10 | 10 |
| Chaetodontidae | *Forcipiger_longirostris* | crustacivore | 7 | 7 |
| Chaetodontidae | *Hemitaurichthys_polylepis* | planktivore | 7 | 3 |
| Chaetodontidae | *Heniochus_chrysostomus* | corallivore | 10 | 10 |
| Chanidae | *Chanos_chanos* | microinvertivore | 1 | 0 |
| Cirrhitidae | *Amblycirrhites_bimacula* | crustacivore | 5 | 0 |
| Cirrhitidae | *Cirrhitichthys_oxycephalus* | crustacivore | 2 | 0 |
| Cirrhitidae | *Cirrhitus_pinnulatus* | crustacivore | 10 | 5 |
| Cirrhitidae | *Neocirrhites_armatus* | crustacivore | 10 | 4 |
| Cirrhitidae | *Paracirrhites_arcatus* | crustacivore | 11 | 12 |
| Cirrhitidae | *Paracirrhites_forsteri* | crustacivore | 10 | 8 |
| Cirrhitidae | *Paracirrhites_hemistictus* | crustacivore | 20 | 12 |
| Creediidae | *Limnichthys_nitidus* | na | 10 | 1 |
| Diodontidae | *Diodon_hysterix* | macroinvertivore | 6 | 6 |
| Ephippidae | *Platax_orbicularis* | na | 2 | 2 |
| Fistulariidae | *Fistularia_commersonii* | piscivore | 10 | 9 |
| Gobiidae | *Asterropteryx_ensifera* | HMD | 1 | 0 |
| Gobiidae | *Asterropteryx_semipunctata* | HMD | 8 | 0 |
| Gobiidae | *Cabillus_tongarevae* | microinvertivore | 9 | 2 |
| Gobiidae | *Callogobius_sclateri* | microinvertivore | 4 | 0 |
| Gobiidae | *Callogobius_sp* | microinvertivore | 6 | 0 |
| Gobiidae | *Ctenogobiops_feroculus* | microinvertivore | 5 | 0 |
| Gobiidae | *Eviota_afelei* | microinvertivore | 14 | 8 |
| Gobiidae | *Eviota_albolineata* | microinvertivore | 17 | 10 |
| Gobiidae | *Eviota_disrupta* | microinvertivore | 2 | 3 |
| Gobiidae | *Eviota_distigma* | microinvertivore | 10 | 6 |
| Gobiidae | *Eviota_infulata* | microinvertivore | 9 | 0 |
| Gobiidae | *Fusigobius_humeralis* | microinvertivore | 4 | 0 |
| Gobiidae | *Fusigobius_neophytus* | microinvertivore | 34 | 11 |
| Gobiidae | *Gnatholepis_cauerensis* | microinvertivore | 30 | 9 |

| | | | | |
|---|---|---|---|---|
| Gobiidae | *Gobiodon_quinquestrigatus* | planktivore | 1 | 0 |
| Gobiidae | *Gobiodon_rivulatus* | planktivore | 2 | 0 |
| Gobiidae | *Nemateleotris_magnifica* | planktivore | 2 | 1 |
| Gobiidae | *Paragobiodon_lacunicolus* | microinvertivore | 10 | 0 |
| Gobiidae | *Paragobiodon_modestus* | microinvertivore | 10 | 3 |
| Gobiidae | *Pleurosicya_labiata* | microinvertivore | 11 | 9 |
| Gobiidae | *Priolepis_ailina* | microinvertivore | 2 | 0 |
| Gobiidae | *Priolepis_compita* | microinvertivore | 12 | 3 |
| Gobiidae | *Priolepis_semidoliata* | microinvertivore | 10 | 11 |
| Gobiidae | *Priolepis_squamogena* | microinvertivore | 10 | 10 |
| Gobiidae | *Priolepis_triops* | microinvertivore | 1 | 0 |
| Gobiidae | *Ptereleotris_evides* | planktivore | 1 | 1 |
| Gobiidae | *Trimmatom_eviotops* | microinvertivore | 9 | 0 |
| Gobiidae | *Valenciennea_strigata* | planktivore | 8 | 4 |
| Hemiramphidae | *Hemiramphus_depauperatus* | piscivore | 4 | 1 |
| Holocentridae | *Myripristis_berndti* | crustacivore | 11 | 3 |
| Holocentridae | *Myripristis_kuntee* | planktivore | 7 | 7 |
| Holocentridae | *Myripristis_violacea* | crustacivore | 10 | 10 |
| Holocentridae | *Neoniphon_argenteus* | crustacivore | 1 | 1 |
| Holocentridae | *Neoniphon_sammara* | crustacivore | 11 | 9 |
| Holocentridae | *Sargocentron_caudimaculatum* | crustacivore | 10 | 2 |
| Holocentridae | *Sargocentron_lepros* | crustacivore | 1 | 0 |
| Holocentridae | *Sargocentron_microstoma* | crustacivore | 5 | 5 |
| Holocentridae | *Sargocentron_spiniferum* | crustacivore | 10 | 10 |
| Holocentridae | *Sargocentron_tiere* | crustacivore | 11 | 10 |
| Kuhlidae | *Kuhlia_mugil* | crustacivore | 4 | 3 |
| Kuhlidae | *Kuhlia_sandvicensis* | crustacivore | 1 | 0 |
| Kyphosidae | *Kyphosus_cinerascens* | HMD | 3 | 3 |
| Kyphosidae | *Kyphosus_vaigiensis* | HMD | 4 | 4 |
| Labridae | *Anampses_caeruleopunctatus* | microinvertivore | 2 | 2 |
| Labridae | *Calotomus_carolinus* | HMD | 10 | 10 |
| Labridae | *Cheilinus_chlorourus* | crustacivore | 11 | 3 |
| Labridae | *Cheilinus_oxycephalus* | crustacivore | 1 | 1 |
| Labridae | *Cheilinus_trilobatus* | macroinvertivore | 7 | 7 |
| Labridae | *Cheilio_inermis* | piscivore | 4 | 2 |
| Labridae | *Chlorurus_microrhinos* | HMD | 6 | 6 |
| Labridae | *Chlorurus_spilurus* | HMD | 11 | 4 |
| Labridae | *Coris_aygula* | macroinvertivore | 10 | 10 |
| Labridae | *Coris_gaimard* | macroinvertivore | 8 | 6 |

| | | | | |
|---|---|---|---|---|
| Labridae | *Epibulus_insidiator* | crustacivore | 10 | 4 |
| Labridae | *Gomphosus_varius* | crustacivore | 10 | 9 |
| Labridae | *Halichoeres_hortulanus* | macroinvertivore | 10 | 9 |
| Labridae | *Halichoeres_marginatus* | microinvertivore | 3 | 3 |
| Labridae | *Halichoeres_trimaculatus* | microinvertivore | 6 | 4 |
| Labridae | *Hemigymnus_fasciatus* | planktivore | 9 | 8 |
| Labridae | *Labroides_bicolor* | microinvertivore | 1 | 0 |
| Labridae | *Novaculichthys_taeniourus* | macroinvertivore | 7 | 7 |
| Labridae | *Oxycheilinus_unifasciatus* | crustacivore | 5 | 4 |
| Labridae | *Pseudocheilinus_hexataenia* | planktivore | 16 | 8 |
| Labridae | *Pseudocheilinus_tetrataenia* | planktivore | 6 | 4 |
| Labridae | *Pseudodax_moluccanus* | macroinvertivore | 2 | 2 |
| Labridae | *Scarus_altipinnis* | HMD | 7 | 4 |
| Labridae | *Scarus_forsteni* | HMD | 11 | 9 |
| Labridae | *Scarus_frenatus* | HMD | 3 | 3 |
| Labridae | *Scarus_globiceps* | HMD | 7 | 7 |
| Labridae | *Scarus_oviceps* | HMD | 9 | 9 |
| Labridae | *Scarus_psittacus* | HMD | 10 | 10 |
| Labridae | *Scarus_rubroviolaceus* | HMD | 5 | 5 |
| Labridae | *Scarus_schlegeli* | HMD | 7 | 7 |
| Labridae | *Thalassoma_hardwicke* | microinvertivore | 10 | 10 |
| Labridae | *Thalassoma_lutescens* | macroinvertivore | 2 | 2 |
| Labridae | *Thalassoma_purpureum* | microinvertivore | 5 | 4 |
| Labridae | *Thalassoma_quinquevittatum* | microinvertivore | 0 | 1 |
| Labridae | *Thalassoma_trilobatum* | microinvertivore | 10 | 3 |
| Lethrinidae | *Gnathodentex_aureolineatus* | macroinvertivore | 10 | 10 |
| Lethrinidae | *Lethrinus_xanthochilus* | piscivore | 2 | 1 |
| Lethrinidae | *Monotaxis_grandoculis* | macroinvertivore | 7 | 6 |
| Lutjanidae | *Aphareus_furca* | piscivore | 10 | 9 |
| Lutjanidae | *Lutjanus_fulvus* | crustacivore | 11 | 11 |
| Lutjanidae | *Lutjanus_kasmira* | crustacivore | 9 | 9 |
| Lutjanidae | *Lutjanus_monostigma* | crustacivore | 5 | 3 |
| Monacanthidae | *Aluterus_scriptus* | corallivore | 4 | 3 |
| Monacanthidae | *Amanses_scopas* | corallivore | 9 | 9 |
| Monacanthidae | *Cantherines_sandwichiensis* | corallivore | 10 | 9 |
| Mugilidae | *Ellochelon_vaigiensis* | HMD | 5 | 5 |
| Mullidae | *Mulloidichthys_flavolineatus* | macroinvertivore | 11 | 10 |
| Mullidae | *Mulloidichthys_vanicolensis* | crustacivore | 5 | 5 |
| Mullidae | *Parupeneus_barberinus* | crustacivore | 4 | 3 |

| | | | | |
|---|---|---|---|---|
| Mullidae | *Parupeneus_ciliatus* | crustacivore | 3 | 3 |
| Mullidae | *Parupeneus_insularis* | crustacivore | 11 | 10 |
| Mullidae | *Parupeneus_multifasciatus* | crustacivore | 11 | 10 |
| Muraenidae | *Gymnothorax_javanicus* | piscivore | 1 | 1 |
| Muraenidae | *Gymnothorax_melatremus* | crustacivore | 7 | 0 |
| Muraenidae | *Uropterygius_alboguttatus* | crustacivore | 6 | 1 |
| Muraenidae | *Uropterygius_fuscoguttatus* | crustacivore | 11 | 7 |
| Ostraciidae | *Ostracion_cubicus* | HMD | 3 | 3 |
| Ostraciidae | *Ostracion_meleagris* | sessile invertivore | 7 | 5 |
| Pempheridae | *Pempheris_oualensis* | microinvertivore | 7 | 3 |
| Pinguipedidae | *Parapercis_millepunctata* | crustacivore | 10 | 8 |
| Platycephalidae | *Onigocia_bimaculata* | microinvertivore | 2 | 0 |
| Pomacanthidae | *Centropyge_bispinosa* | sessile invertivore | 10 | 8 |
| Pomacanthidae | *Centropyge_flavissima* | sessile invertivore | 10 | 7 |
| Pomacanthidae | *Centropyge_loricula* | sessile invertivore | 5 | 3 |
| Pomacanthidae | *Pomacanthus_imperator* | sessile invertivore | 7 | 5 |
| Pomacanthidae | *Pygoplites_diacanthus* | sessile invertivore | 11 | 10 |
| Pomacentridae | *Abudefduf_septemfasciatus* | planktivore | 10 | 9 |
| Pomacentridae | *Abudefduf_sexfasciatus* | planktivore | 10 | 2 |
| Pomacentridae | *Abudefduf_sordidus* | microinvertivore | 6 | 4 |
| Pomacentridae | *Chromis_acares* | planktivore | 8 | 8 |
| Pomacentridae | *Chromis_atripectoralis* | planktivore | 11 | 12 |
| Pomacentridae | *Chromis_iomelas* | planktivore | 11 | 8 |
| Pomacentridae | *Chromis_margaritifer* | planktivore | 10 | 12 |
| Pomacentridae | *Chromis_vanderbilti* | planktivore | 7 | 7 |
| Pomacentridae | *Chromis_xanthura* | planktivore | 10 | 6 |
| Pomacentridae | *Chrysiptera_brownriggii* | HMD | 10 | 1 |
| Pomacentridae | *Dascyllus_aruanus* | planktivore | 10 | 6 |
| Pomacentridae | *Dascyllus_flavicaudus* | planktivore | 11 | 10 |
| Pomacentridae | *Dascyllus_trimaculatus* | planktivore | 10 | 9 |
| Pomacentridae | *Plectroglyphidodon_dickii* | HMD | 4 | 2 |
| Pomacentridae | *Plectroglyphidodon_johnstonianus* | corallivore | 10 | 2 |
| Pomacentridae | *Plectroglyphidodon_lacrymatus* | HMD | 7 | 0 |
| Pomacentridae | *Pomacentrus_coelestis* | planktivore | 1 | 1 |
| Pomacentridae | *Pomacentrus_pavo* | planktivore | 10 | 2 |
| Pomacentridae | *Pomachromis_fuscidorsalis* | planktivore | 1 | 1 |
| Pomacentridae | *Stegastes_fasciolatus* | HMD | 10 | 8 |
| Pomacentridae | *Stegastes_nigricans* | HMD | 11 | 2 |
| Priacanthidae | *Priacanthus_hamrur* | crustacivore | 1 | 1 |

| | | | | |
|---|---|---|---|---|
| Pseudochromidae | *Pseudoplesiops_revellei* | microinvertivore | 5 | 0 |
| Scombridae | *Gymnosarda_unicolor* | piscivore | 1 | 1 |
| Scorpaenidae | *Caracanthus_maculatus* | crustacivore | 10 | 5 |
| Scorpaenidae | *Caracanthus_unipinna* | crustacivore | 6 | 0 |
| Scorpaenidae | *Pterois_radiata* | crustacivore | 10 | 9 |
| Scorpaenidae | *Scorpaenodes_parvipinnis* | crustacivore | 4 | 0 |
| Scorpaenidae | *Scorpaenodes_scaber* | crustacivore | 10 | 8 |
| Scorpaenidae | *Scorpaenodes_varipinnis* | crustacivore | 1 | 0 |
| Scorpaenidae | *Scorpaenopsis_diabolus* | crustacivore | 1 | 0 |
| Scorpaenidae | *Scorpaenopsis_papuensis* | crustacivore | 1 | 1 |
| Scorpaenidae | *Sebastapistes_fowleri* | crustacivore | 10 | 13 |
| Scorpaenidae | *Sebastapistes_mauritiana* | crustacivore | 10 | 0 |
| Serranidae | *Cephalopholis_argus* | piscivore | 12 | 4 |
| Serranidae | *Cephalopholis_aurantia* | crustacivore | 1 | 0 |
| Serranidae | *Cephalopholis_sexmaculata* | piscivore | 2 | 1 |
| Serranidae | *Cephalopholis_urodeta* | crustacivore | 10 | 9 |
| Serranidae | *Epinephelus_fasciatus* | crustacivore | 6 | 4 |
| Serranidae | *Epinephelus_hexagonatus* | piscivore | 10 | 9 |
| Serranidae | *Epinephelus_merra* | crustacivore | 11 | 5 |
| Serranidae | *Plectranthias_nanus* | crustacivore | 11 | 9 |
| Serranidae | *Pseudanthias_pascalus* | crustacivore | 7 | 7 |
| Serranidae | *Pseudogramma_polyacanthum* | crustacivore | 20 | 4 |
| Serranidae | *Variola_louti* | piscivore | 8 | 2 |
| Siganidae | *Siganus_spinus* | HMD | 5 | 5 |
| Sphyraenidae | *Sphyraena_barracuda* | piscivore | 2 | 2 |
| Syngnathidae | *Corythoichthys_flavofasciatus* | planktivore | 4 | 4 |
| Syngnathidae | *Doryrhamphus_excisus* | planktivore | 2 | 1 |
| Synodontidae | *Synodus_binotatus* | piscivore | 4 | 3 |
| Tetradontidae | *Arothron_meleagris* | corallivore | 10 | 9 |
| Tetradontidae | *Canthigaster_bennetti* | sessile invertivore | 3 | 3 |
| Tetradontidae | *Canthigaster_janthinoptera* | sessile invertivore | 5 | 2 |
| Tetradontidae | *Canthigaster_solandri* | HMD | 10 | 6 |
| Tetradontidae | *Canthigaster_valentini* | sessile invertivore | 3 | 2 |
| Tripterygiidae | *Enneapterygius_pyramis* | microinvertivore | 24 | 14 |
| Tripterygiidae | *Enneapterygius_sp* | microinvertivore | 11 | 0 |
| Zanclidae | *Zanclus_cornutus* | sessile invertivore | 10 | 10 |
| **TOTAL** | | | **1938** | **1270** |

**Table S2.**

The 18S, COI, and 23S primer sets used for gut content DNA metabarcoding.

| Gene region | Primer | Sequence 5'-3' | Reference |
|---|---|---|---|
| 18S | V4_18SF | CCAGCASCYGCGGTAATTCC | Piredda et al. (2017) |
| | V4_18SR | ACTTTCGTTCTTGATYRATGA | |
| COI | mICOIintF | GGWACWGGWTGAACWGTWTAYCCYCC | Leray et al. (2013) |
| | jgHCO2198 | TAIACYTCIGGRTGICCRAARAAYCA | Geller et al. (2013) |
| 23S | p23SrV_fl | GGACAGAAAGACCCTATGAA | Sherwood and Presting (2007) |
| | DIam23Sr1 | TGAGTGACGGCCTTTCCACT | |

**Table S3.**

Prey item categories for the trophic network plot (Fig. 2). Prey items were grouped into six categories, with categories assigned based on prey phylum, except Arthropoda, which was assigned based on prey class ("na" indicates that class was not used for categorization).

| Prey phylum | Prey class | Network category |
|---|---|---|
| Acanthocephala | na | Microinvertebrates |
| Annelida | na | Macroinvertebrates |
| Arthropoda | Arachnida | Macroinvertebrates |
| Arthropoda | Branchiopoda | Macroinvertebrates |
| Arthropoda | Copepoda | Microinvertebrates |
| Arthropoda | Hexapoda | Macroinvertebrates |
| Arthropoda | Insecta | Macroinvertebrates |
| Arthropoda | Malacostraca | Macroinvertebrates |
| Arthropoda | Ostracoda | Microinvertebrates |
| Arthropoda | Pycnogonida | Macroinvertebrates |
| Arthropoda | Thecostraca | Sessile Invertebrates |
| Bigyra | na | Microbes |
| Bryozoa | na | Sessile Invertebrates |
| Cercozoa | na | Microbes |
| Chaetognatha | na | Microinvertebrates |
| Chlorophyta | na | Algae |
| Chordata | na | Vertebrates |
| Ciliophora | na | Microbes |
| Cnidaria | na | Sessile Invertebrates |
| Cryptophyta | na | Microbes |
| Ctenophora | na | Macroinvertebrates |
| Cyanobacteria | na | Microbes |
| Echinodermata | na | Macroinvertebrates |
| Entoprocta | na | Sessile Invertebrates |
| Euglenozoa | na | Microbes |
| Gastrotricha | na | Microinvertebrates |
| Haptista | na | Microbes |
| Haptophyta | na | Microbes |
| Marchantiophyta | na | Algae |
| Mollusca | na | Macroinvertebrates |
| Myzozoa | na | Microbes |
| Nematoda | na | Microinvertebrates |
| Nemertea | na | Microinvertebrates |

| | | |
|---|---|---|
| Ochrophyta | na | Algae |
| Oomycota | na | Microbes |
| Phoronida | na | Sessile Invertebrates |
| Phragmoplastophyta | na | Algae |
| Platyhelminthes | na | Microinvertebrates |
| Porifera | na | Sessile Invertebrates |
| Retaria | na | Microbes |
| Rhodophyta | na | Algae |
| Rotifera | na | Microinvertebrates |
| Tracheophyta | na | Algae |
| Xenacoelomorpha | na | Microinvertebrates |

**Table S4.**

Collinearity diagnostics across model formulations. Maximum condition index and maximum absolute Pearson correlation between richness and niche segregation from models including richness, segregation, their interaction, and reef habitat, for each dataset (metabarcoding and isotope) and weighting scheme (weighted and unweighted). Condition index values below 30 indicate the absence of severe multicollinearity.

| Model | Max condition index | Max Pearson correlation |
|---|---|---|
| Metabarcoding weighted | 23.203 | 0.789 |
| Metabarcoding unweighted | 4.749 | 0.078 |
| Isotope weighted | 13.764 | 0.586 |
| Isotope unweighted | 4.539 | 0.287 |

**Table S5.**

Diagnostics and predictive performance for Bayesian models of niche segregation. For each model (meta = metabarcoding; iso = isotope; w = weighted; unw = unweighted; weak = weakly informative priors; noint = no interaction term), we report the number of observations (n_obs), the number and fraction of divergent transitions (n_diverg, frac_diverg), the maximum tree depth reached during sampling (max_treedepth), the maximum Gelman–Rubin statistic across parameters (max_Rhat), and the minimum bulk and tail effective sample sizes (min_bulkESS, min_tailESS). Predictive performance is assessed using leave-one-out (LOO) cross-validation, reported as the LOO information criterion (looic) and expected log predictive density (elpd_loo). Overall model fit is summarized by Bayesian $R^2$ (bayes_$R^2$). All models showed satisfactory convergence and sampling efficiency.

| model | n_obs | n_diverg | frac_diverg | max_treedepth | max_Rhat | min_bulkESS | min_tailESS | looic | elpd_loo | bayes_$R^2$ |
|---|---|---|---|---|---|---|---|---|---|---|
| meta_unw_weak | 2172 | 0 | 0 | 6 | 1.000 | 11037 | 10423 | 4130 | -2065 | 0.46 |
| meta_unw_weak_noint | 2172 | 0 | 0 | 5 | 1.000 | 9528 | 9777 | 4138 | -2069 | 0.45 |
| meta_w_weak | 2172 | 0 | 0 | 7 | 1.000 | 7058 | 9477 | 4225 | -2112 | 0.44 |
| meta_w_weak_noint | 2172 | 0 | 0 | 6 | 1.000 | 10614 | 9917 | 4276 | -2138 | 0.43 |
| iso_unw_weak | 2131 | 0 | 0 | 6 | 1.000 | 9430 | 10398 | 3635 | -1817 | 0.51 |
| iso_w_weak | 2131 | 0 | 0 | 7 | 1.000 | 8961 | 9249 | 3651 | -1825 | 0.50 |
| iso_unw_weak_noint | 2131 | 0 | 0 | 5 | 1.000 | 10123 | 10531 | 3661 | -1830 | 0.50 |
| iso_w_weak_noint | 2131 | 0 | 0 | 6 | 1.001 | 11474 | 10413 | 3735 | -1867 | 0.49 |

**Data S1. (separate file)**

Metadata for sequencing libraries deposited on NCBI under BioProject PRJNA1522544. Data include Sequence Read Archive (SRA) run accessions (sra_accession), BioSample accessions (biosample_accession), sample names (sample_name), fish host species (fish_species), and paired-end FASTQ filenames for forward (R1; filename1) and reverse (R2; filename2) reads for all fish gut content metabarcoding libraries generated in this study. Sequencing libraries with zero reads were not submitted to NCBI, so the SRA run accessions are marked as NA. The final column (status) indicates whether the library yielded sequence data ("Data available") or was excluded due to zero-read files ("No data, removed"). Of the 5,097 paired sequencing libraries, 63 libraries had zero reads, which were distributed across 39 fish samples.
